# Axon guidance at the midline – a live imaging perspective

**DOI:** 10.1101/2020.07.20.211995

**Authors:** Alexandre Dumoulin, Nikole R. Zuñiga, Esther T. Stoeckli

## Abstract

During neural circuit formation, axons navigate several choice points to reach their final target. At each one of these intermediate targets, growth cones need to switch responsiveness from attraction to repulsion in order to move on. Molecular mechanisms that allow for the precise timing of surface expression of a new set of receptors that support the switch in responsiveness are difficult to study *in vivo*. Mostly, mechanisms are inferred from the observation of snapshots of many different growth cones analyzed in different preparations of tissue harvested at distinct time points. However, to really understand the behavior of growth cones at choice points, a single growth cone should be followed arriving at and leaving the intermediate target.

Here, we describe a spinal cord preparation that allows for live imaging of individual axons during navigation in their intact environment. The possibility to observe single growth cones navigating their intermediate target allows for measuring growth speed, changes in morphology, or aberrant behavior. Moreover, observation of the intermediate target – the floor plate – revealed its active participation and interaction with commissural axons during midline crossing.

**Summary statement:** Live tracking of single growth cones is more informative about axonal behavior during navigation than inference of behavior from the analyses of snapshots of different growth cones.

## INTRODUCTION

Commissural axons in the developing spinal cord have been used for over two decades to learn fundamental molecular mechanisms of axon guidance (Stoeckli, 2018). The focus was on the dI1 subtype of dorsal commissural interneurons, as their axons have a very stereotypical trajectory at the ventral midline, where they all cross the floor plate (FP), exit it and turn rostrally along the contralateral border. Thus, dI1 commissural neurons offer an easy read-out for deciphering molecular mechanisms of axon guidance at choice points. Since the first application of lipophilic dye tracing in open-book preparations of rat spinal cords that revealed the normal trajectory of these axons 30 years ago (Bovolenta and Dodd, 1990), this method continues to be used to assess axon guidance at the midline in mouse and chicken embryos. The comparison between axons in open-book preparations of control and experimentally manipulated spinal cords, dissected at specific time points, offered a solid understanding of molecules involved in axonal midline crossing and subsequent turning in higher vertebrates.

However, the information about mechanisms that can be extracted from such experiments is limited, as it is deduced from snapshots of axons taken from different animals. For this reason, we have established a live-imaging approach that allows for visualization of axonal behavior while they are crossing the midline and then turning rostrally. We have chosen the chicken embryo, as it is a very accessible model for studying various developmental processes in intact tissues *in vivo* and *ex vivo* (Sanders et al., 2013; Das and Storey, 2014; Boubakar et al., 2017; Li et al., 2019). Thanks to a very stable and reproducible spinal cord culture, we could for the first time characterize the exact timing of midline crossing and the details of rostral turning by dI1 axons in an intact environment in control and experimentally manipulated spinal cords. We could get more insight into growth cone dynamics and morphologies at choice points. Furthermore, our *ex vivo* method also shed new light on the role of the intermediate target, the FP cells, their dynamics, morphology and interaction with commissural axons during midline crossing.

## RESULTS

### Electroporation is an efficient tool to selectively label dI1 neurons

We used unilateral *in ovo* electroporation of the chicken spinal cord in Hamburger and Hamilton (HH) stage 17-18 embryos to specifically express farnesylated td-Tomato (td-Tomato-F) in Math1-positive dI1 neurons, as well as farnesylated EGFP (EGFP-F) expression in their environment (Fig. 1A,B). One day after electroporation, at HH22, embryos showed expression of td-Tomato-F restricted to dI1 neurons and EGFP-F expression in the entire half of the spinal cord, as expected (Fig. 1C) (Wilson and Stoeckli, 2011). At this stage, most of the Math1-positive dI1 axons approached the FP, but did not yet cross the midline, whereas other more ventral populations of commissural neurons expressing EGFP-F already projected many axons to the contralateral side of the spinal cord (white arrowheads and arrow, respectively, Fig. 1C). For this reason, we chose HH22 as the optimal stage to start tracing dI1 axons at the midline using live imaging.

**Fig. 1.**
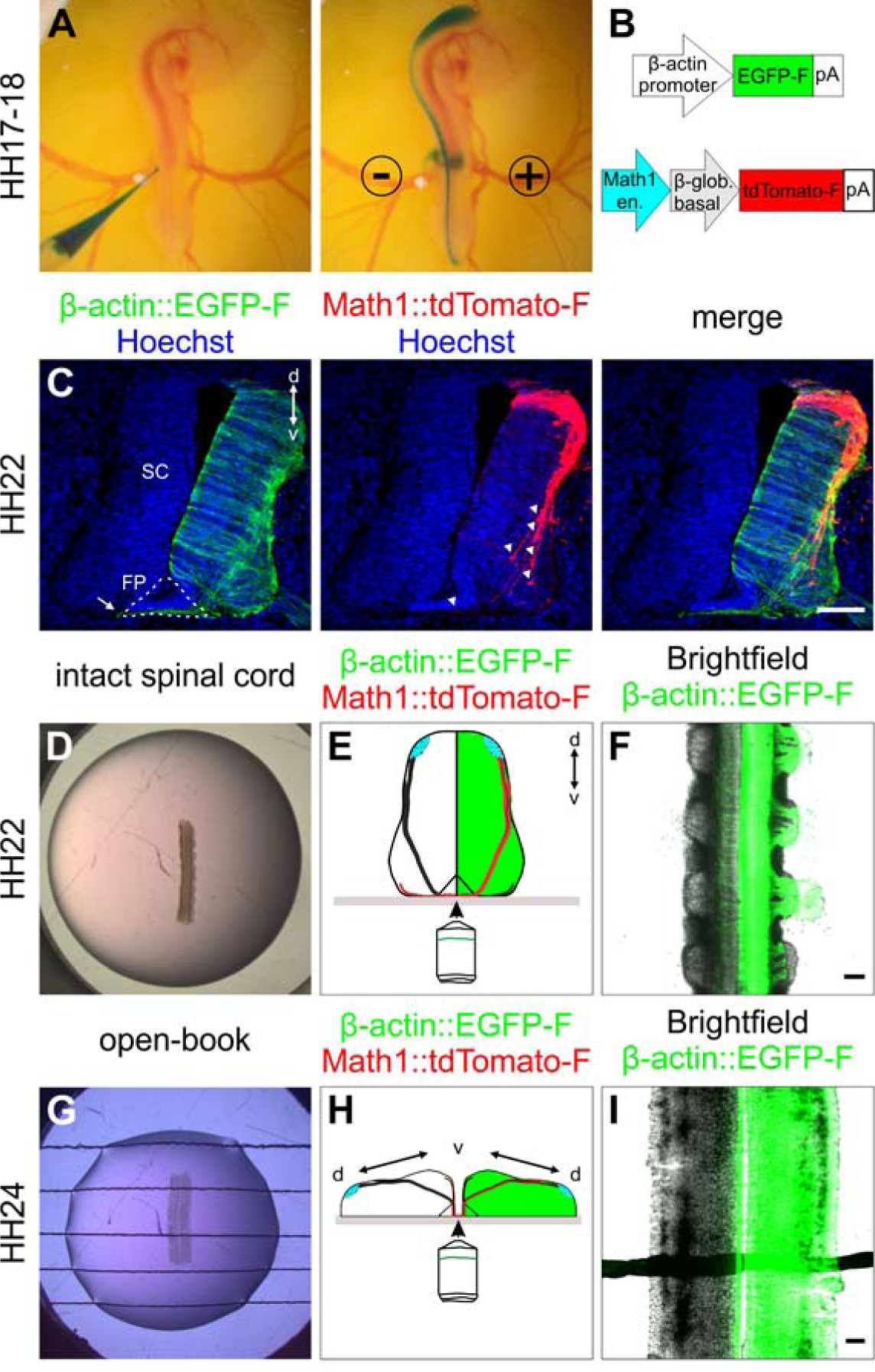
Labeling strategy for dI1 interneurons and spinal cord culture systems. (A-C) *In ovo* injection and electroporation of a plasmid mix to specifically label dI1 interneurons. (A) The plasmid mix was injected into the central canal of the spinal cord of HH17-18 chicken embryo *in ovo*, followed by unilateral electroporation. (B) Plasmid constructs injected to target all cells (β-actin::EGFP-F) and dI1 interneurons β (Math1::tdTomato-F). en., enhancer; β-glob., β-globin. (C) Immunostaining of a transverse cryosection of a HH22 spinal cord taken from an embryo sacrificed one day after electroporation with the plasmids indicated in (B). At this stage, most dI1 growth cones were approaching the FP area, but none of them had crossed it yet (white arrowheads). However, a substantial number of Math1-negative, but EGFP-F-expressing commissural axons of more ventral populations had already crossed the FP at HH22 (arrow). (D-F) Intact spinal cord culture. (D) Intact spinal cords of embryos injected and electroporated one day earlier were embedded with the ventral side down in a drop of low-melting agarose. (E) The ventral spinal cord area was imaged with an inverted spinning disk microscope. The green-colored hemisphere represents the electroporated side of the spinal cord. (F) Low magnification overview of a spinal cord visualized with this set-up with cells expressing EGFP-F under the β-actin promoter on one side merged with the brightfield image. (G-I) Culture of an open-book preparation of a spinal cord. (G) Intact open-book preparations of HH24 spinal cords dissected from embryos injected and electroporated about one and a half day earlier were embedded with the apical side down in an agarose drop with strings to hold it in place. (H) The midline area was visualized with the same inverted spinning disk microscope as above. The green-colored hemisphere represents the electroporated side of the spinal cord. (I) Low magnification overview of a spinal cord visualized with this set-up with cells expressing EGFP-F under the β-actin promoter on one side merged with the brightfield image. SC, spinal cord; d, dorsal; v, ventral. Scale bars: 100 µm.

### Live imaging of dI1 axons at the midline of intact spinal cord

We extracted the intact spinal cord one day after electroporation (Fig. S1), cultured it with the ventral midline down and imaged it with an inverted spinning disk microscope and a 20x objective (Fig. 1D-F). Math1-positive dI1 axons crossing the FP could be visualized for at least 24 hours (white arrowheads, Fig. 2A; Movies 1, 2). Within this time window many dI1 axons crossed the midline, exited the FP, turned rostrally and formed the contralateral ventral funiculus (white arrows, Fig. 2A). A Math1-positive ipsilateral subpopulation of axons could also be seen in these recordings as previously reported *in vivo* (Phan et al., 2010) (white asterisk, Fig. 2A). Cultures of intact spinal cords turned out to be a very stable system as the U-shaped morphology of the commissure was preserved over time (Fig. 2B, Movie 2) and all major cell populations were still in place after one day *ex vivo* (Fig. S2). Furthermore, Sonic hedgehog (Shh) expression was still restricted to the FP and showed the caudal (high) to rostral (low) gradient like *in vivo* (Fig. S3) (Bourikas et al., 2005), and most importantly, dI1 axons’ navigation was identical to the *in vivo* situation during this time window (Fig. S4). We also compared our *ex vivo* method with a recently published protocol using open-book preparations of HH24-26 chicken spinal cords (Pignata et al., 2019) (Fig. 1G-I). In contrast to this protocol, our *ex vivo* method did not result in overshooting axons, an artefact that was already seen after short times in cultures of open-book preparations, with Math1-positive dI1 commissural axons that crossed the midline but then failed to turn into the longitudinal axis and continued to grow straight into the contralateral side instead (white arrowheads, Fig. 2C,D; Movie 3). Although open-book cultures offer the possibility to follow midline crossing of dI1 axons, the deformation of the FP and commissure in this preparation (Fig. 2D) and the fact that diffusible guidance cues are not well retained in the tissue, most likely lead to these artifacts. These problems were not seen in our cultures of intact spinal cords with meninges attached that prevents diffusion of secreted molecules and preserves gradients (Fig. S3). Therefore, our *ex vivo* method of culturing intact spinal cords offers a highly stable intact system in which dI1 commissural axons are behaving as expected based on what is known from *in vivo* studies.

**Fig. 2.**
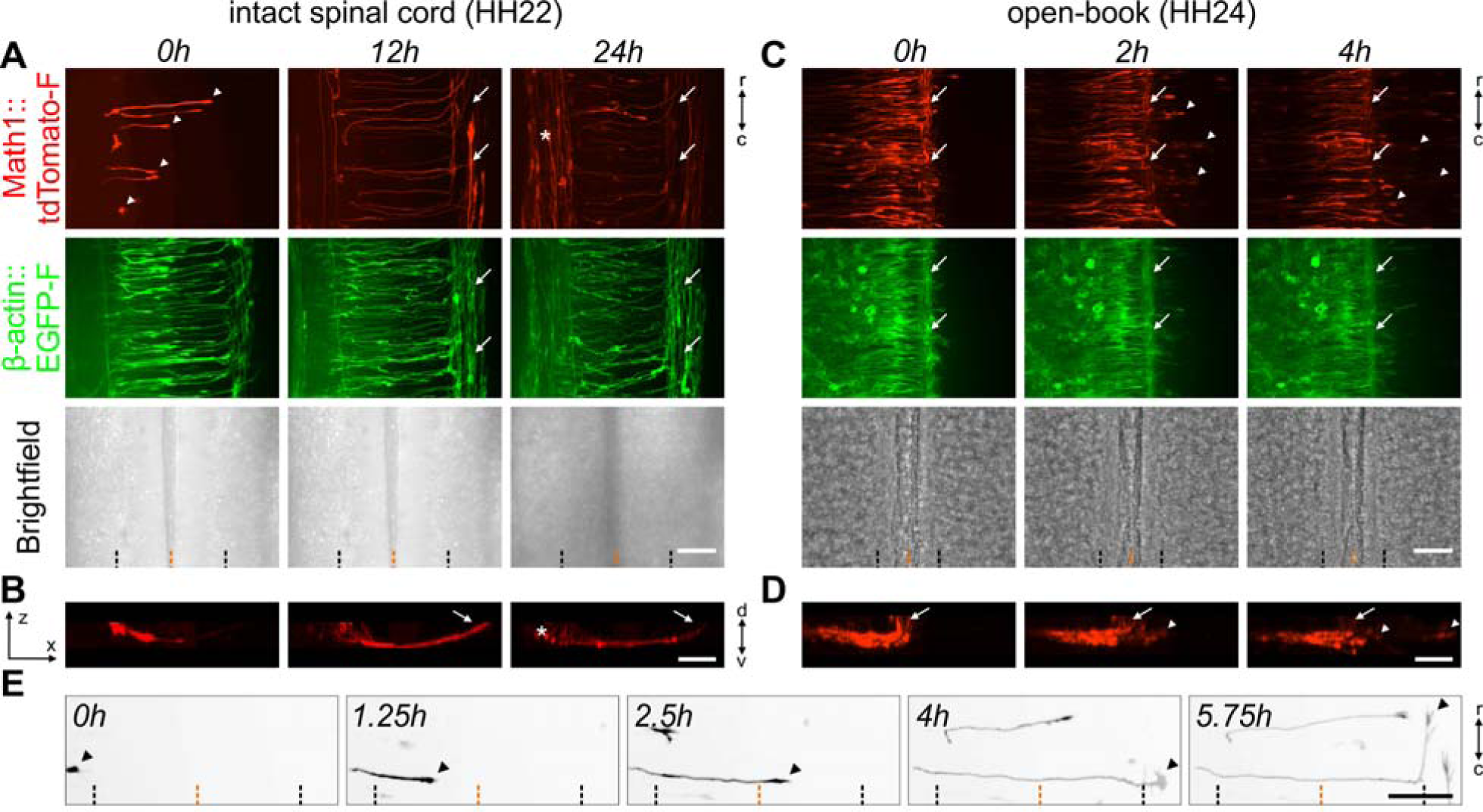
Live imaging of cultured intact spinal cords allowed for the visualization of dI1 axons during floor-plate crossing and navigation into the longitudinal axis. (A) 24-h time-lapse recording showed that Math1-positive dI1 commissural axons could cross the FP (white arrowheads), turn anteriorly, as expected, and form the contralateral ventral funiculus (white arrows) in cultured intact HH22 spinal cords. The asterisk indicates a population of Math1-positive ipsilateral axons. (B) Transversal view of a region of interest from the time-lapse recording shown in (A), highlighting the trajectory of dI1 axons and the formation of the commissure. The white arrow indicates the position of the contralateral ventral funiculus. The white asterisk labels ipsilateral axons. (C) 4-h time-lapse recording showing Math1-positive dI1 commissural axons in a cultured open-book preparation of a HH24 spinal cord. Note that within less than 2h in culture the majority of dI1 commissural axons were overshooting the contralateral FP boundary and growing straight into the contralateral side after having crossed the FP (white arrowheads). White arrows indicate the contralateral ventral funiculus. (D) Transversal view of a region of interest from the time-lapse sequence shown in (C) highlighting the aberrant trajectory of dI1 commissural axons (white arrowheads) growing straight past the contralateral ventral funiculus (white arrow). (E) Single dI1 growth cones (black arrowheads) could be tracked crossing the FP, exiting it and turning rostrally. Math1-positive axons are now shown in black. Black and orange dashed lines indicate FP boundaries and the midline, respectively. d, dorsal; v, ventral; r, rostral; c, caudal. Scale bars: 50 µm.

### Characterization of the timing of midline crossing by dI1 commissural axons

The time it takes commissural axons to cross the FP has been estimated but could not be measured exactly (Stoeckli, 2018; Zou, 2012). However, timing is an issue, because axons have to change their responsiveness to FP-derived guidance cues, like the Slits, Shh, or Wnt proteins, by expressing appropriate receptors in a precisely regulated manner (Bourikas et al., 2005; Domanitskaya et al., 2010; Long et al., 2004; Lyuksyutova et al., 2003; Philipp et al., 2012; Wilson and Stoeckli, 2013). With our method, we could track single dI1 commissural axons in the FP at any time point and in different regions of interest (black arrowheads, Fig. 2E). We were therefore able to ask, how long dI1 axons needed for FP crossing and their subsequent rostral turn (Fig. 3A). On average, dI1 commissural axons took 5.6±1.4h to cross the entire FP and 1.4±1.0h to turn and initiate the rostral growth at the FP exit site. Thus, in total, they needed 6.9±1.8h from entering the FP to the initiation of their rostral growth (mean ± standard deviation, Fig. 3B and Table S1). There was no significant difference between the average time of crossing the first versus the second half of the FP (Fig. 3B, Table S1).

**Fig. 3.**
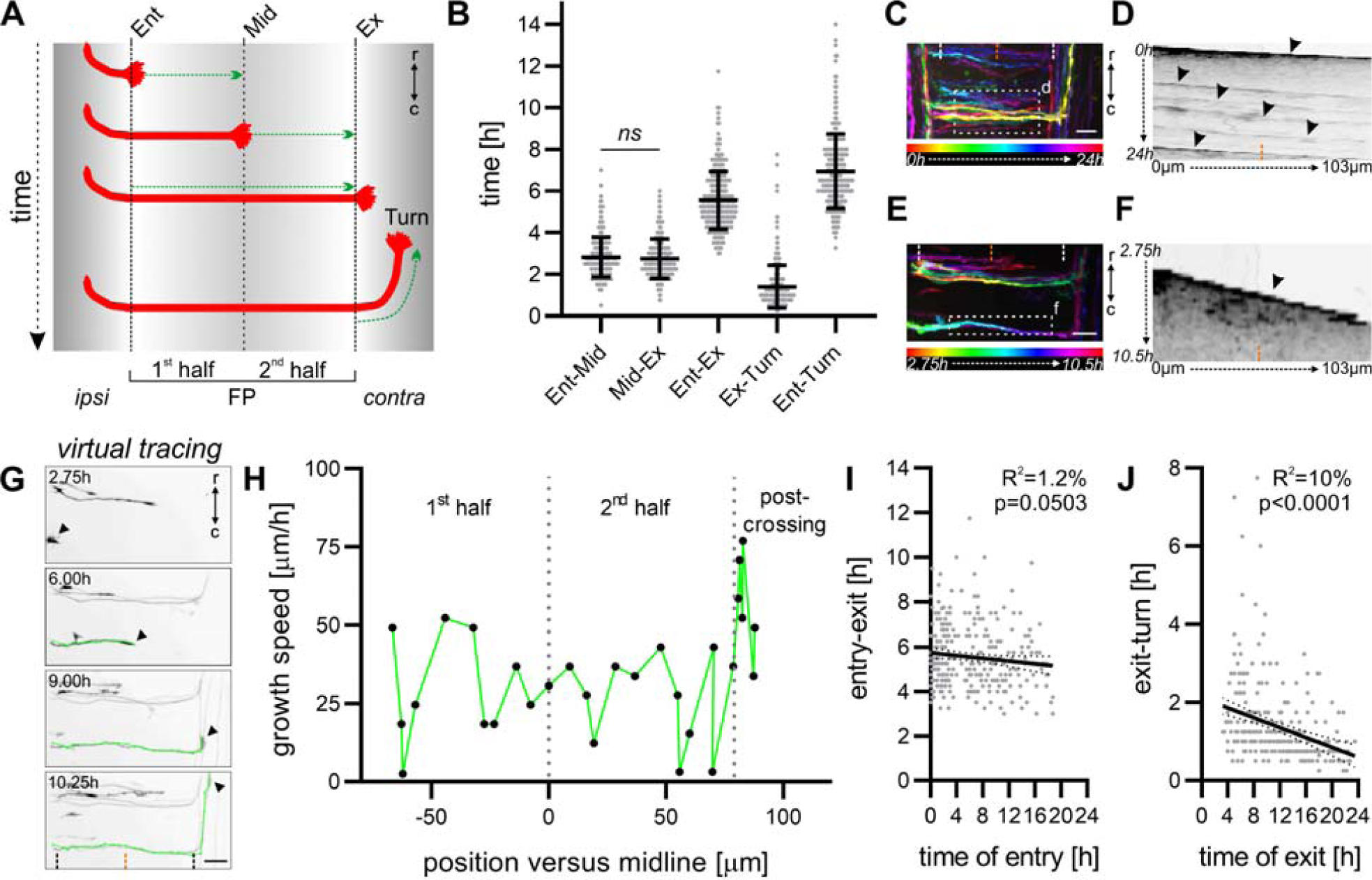
Characterization of the timing of midline crossing by dI1 commissural axons. (A) Schematic depicting how the timing of midline crossing for individual dI1 axons was measured for each segment of interest. (B) Graph showing the average time a growth cone spent in each segment shown in (A) (N_embryos_=7; n_axons_= 298). There was no significant difference in the time taken by dI1 axons to cross the first versus the second half of the FP (p≥0.05, paired two-tailed Wilcoxon test). (C) Temporal-color code projection of a 24-h time-lapse recording. (D) Kymograph of the 24-h time-lapse recording in the region of interest within the FP shown in (C). Several axons crossing the FP at different times can be visualized (black arrowheads). (E) Temporal-color code projection of 7.75-h time-lapse recording segment. (F) Kymograph of the 7.75-h time-lapse recording segment in the region of interest within the FP shown in (E). Within this time segment only one axon crossed the FP and could be visualized in the kymograph (black arrowhead). White and orange dashed lines represent the FP boundaries and the midline, respectively. (G) A virtual tracing tool (shown in green) was used to extract the velocity of the growth cones (black arrowheads) at each time point during midline crossing at the single axon level. The same axon is shown in (E) and (G). Black dashed lines and the orange dashed lines represent the FP boundaries and midline, respectively. (H) The growth speed of the axon shown in (G) could be extracted and plotted against the position of the growth cone in the FP. Dotted grey lines represent the time at which the axon crossed the midline or exited the FP. (I,J) The time of crossing the FP (entry-exit) or of turning (exit-turn) for each axons measured in (B) was plotted against the time of FP entry and exit of the growth cone, respectively. (I) The time axons took to cross the FP appeared to decrease over time although this was not significant. (J) The time axons took to turn after exiting the FP decreased significantly over time. ipsi, ipsilateral; contra, contralateral; Ent, entry; Mid, midline; Ex, exit; r, rostral; c, caudal. Scale bars: 25 µm.

This was supported by kymographic analysis of a region of interest within the FP from 24-hour time-lapse recordings showing similar growth patterns between the first and second half of the FP at the single axon level (Fig. 3C, black arrowheads in Fig. 3D, Movie 4). Although the kymographic analysis was useful to screen for overall growth pattern of single dI1 axons within the FP (Fig. 3E,F), it was not sensitive enough to detect more subtle changes in growth speed. Hence, we used a virtual tracing tool to follow the movement of the leading edge of each growth cone at each time point (Fig. 3G). With this tool we could extract the instantaneous growth speed for each axon. It turned out that the large majority of them had a fluctuating growth pattern with random acceleration-deceleration pulses that could be observed in early as well as late crossing axons (Fig. 3H, Movie 5, Fig. S5). Another interesting observation was made when we compared the times of crossing the FP and the initiation of rostral growth after turning. It seemed that there was a trend towards reduced time of FP crossing in later crossing axons, but this was not significant (p=0.0503; Fig.3I). However, the time dI1 axons took to turn rostrally at the exit site was significantly reduced over time (Fig. 3J). The latter observation suggest that commissural axons that already turned anteriorly at the contralateral FP exit site might help the following ones to turn more rapidly. Our method offers new opportunities for further investigations of possible collaborations between axons at choice points.

### Characterization of the dI1 growth cone morphology at choice points

Another aspect that we considered was growth cone morphology. The growth cone plays a central role in axon guidance, as it explores the environment for guidance cues and translates this information into the directionality of growth (Stoeckli, 2018; de Ramon Francàs et al., 2017). We observed that dI1 growth cones in the FP appeared to have a thin and elongated shape in the direction of growth. At the FP exit site, they transiently enlarged (arrowheads, Fig. 4D, Movie 2). We measured the average growth cone area in each segment of interest and confirmed that growth cones at the exit site of the FP were indeed significantly larger than the ones within the FP or after the turn (Fig. 4A,B, Table S1). There was no significant change in the average growth cone area between the first and second half of the FP (Fig. 4B, Table S1). The changes of growth cone shape were in line with previous reports on chicken and rat commissural axons *in vivo* (Bovolenta and Dodd, 1990; Yaginuma et al., 1991) and our data on Math1-positive axons *in vivo* (Fig. 4C, Table S1, Fig. S4B). The possibility to follow individual axons over time allowed us to make novel observations of their behavior at the FP exit site. The growth cones very often extended long filopodia in both rostral and caudal direction just before turning (Fig. 4D, Movies 6, 7). Some of the growth cones even appeared to transiently split just before turning rostrally, similar to dorsal root ganglia central afferents in the mouse dorsal root entry zone before bifurcating (Dumoulin et al., 2018) (Movie 8). All these features are present *in vivo*, as similar growth cone morphologies were found in fixed HH24-25 spinal cords (Fig. 4E). Thus, *ex vivo* live imaging of cultured intact spinal cords using low magnification time-lapse microscopy offers the opportunity to detect morphological changes of growth cones at choice points and is ideal for the analysis of many aspects of midline crossing. However, the limited resolution especially in 3D might preclude the detection of more subtle changes in morphology of growth cones while crossing the midline.

**Fig. 4.**
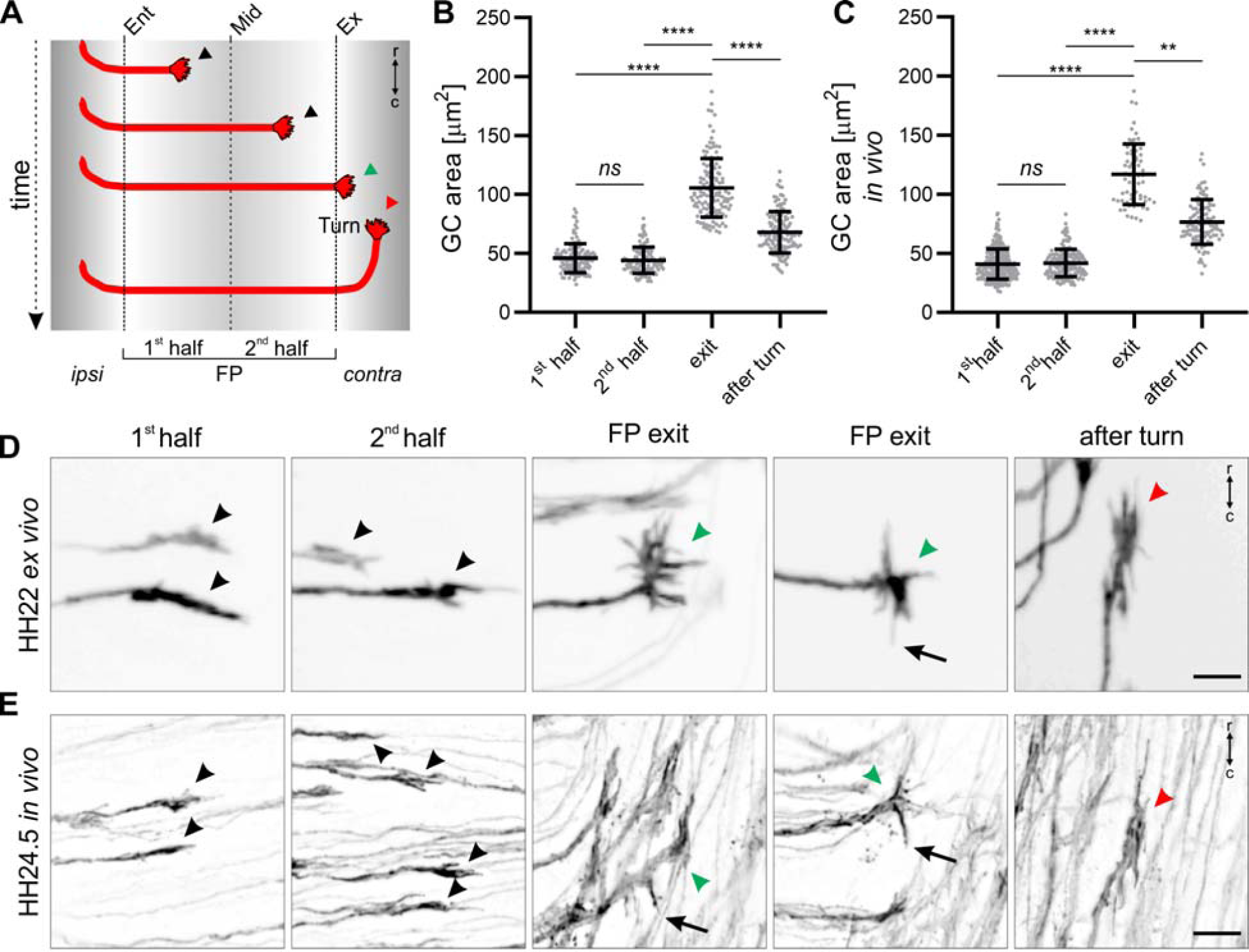
Live imaging of intact spinal cords revealed dI1 growth cone morphologies at chosen time points. (A) Schematic depicting where the growth cone area of individual dI1 axons was measured for (B) and (C). (B) Average growth cone areas were measured from 24-h time-lapse recordings of dI1 axons crossing the midline (N_embryos_=7; n_growth cones_= 127). No significant difference in the area of growth cones was found between the first and second half of the FP. However, growth cones were significantly larger at the exit site but then again reduced in size after having turned rostrally (paired Friedman test with Dunn’s multiple-comparisons test). (C) Average growth cone areas were measured *in vivo* from fixed HH23-25 spinal cords (N_embryos_=8; n_growth cones_= 285(1^st^ half), 153 (2^nd^ half), 68 (exit) and 102 (after turn). The relationship between the average growth cone area and the position in the FP corroborated results using the *ex vivo* culture system shown in (B) (unpaired Kruskal-Wallis test with Dunn’s multiple-comparisons test). (D,E) Examples corroborating the similarities in growth cone morphology *ex vivo* and *in vivo* in the FP (black arrowheads), at the exit site (green arrowheads) and after rostral turn (red arrowheads). At the exit site, growth cones were spiky with always some filopodia pointing caudally just before rostral turn, a feature that was also observed *in vivo* (black arrows). ipsi, ipsilateral; contra, contralateral; r, rostral; c, caudal. Error bars represent standard deviation. p<0.0001 (****), p<0.01 (**) and p≥0.05 (ns) for all tests. Scale bars: 10 µm.

### Higher magnification analysis of commissural axons crossing the midline revealed dorso-ventral activities of their growth cone

For this reason, we repeated time-lapse recordings of cultured intact spinal cords using a higher magnification objective and advanced 3D deconvolution technology. This allowed following Math1::tdTomato-F-positive dI1 axons over time while entering, crossing and exiting the FP (Movie 9). We observed that dI1 growth cones crossing the FP were bulkier in the dorso-ventral axis (white arrowheads, Fig. 5A, Movie 10) and were particularly dynamic in this axis showing rapid extension of filopodial protrusions (white arrows in Fig. 5A and Movie 10). This aspect of dI1 growth cone behavior in the FP was supported by immunostaining of HH22-24.5 whole-mount spinal cords or cryosections (arrowheads, Fig. 5B-D, Movie 11,12). In line with their dorso-ventral activity we could observe that dI1 growth cones in the commissure sent a dynamic long protrusion (up to ∼13 μm) into the FP while crossing it (black arrow, Fig. 5E, black arrowheads, Movie 13). Protrusions entering the FP could also be detected for dI1 growth cones *in vivo* as revealed by immunostaining of HH22-24.5 whole-mount spinal cords or cryosections (white arrows, Fig. 5F,G). Next, we also had a closer look at the FP exit site, where growth cones need to read longitudinal gradients to initiate the rostral turn after exiting the FP (Pignata et al., 2019; Stoeckli, 2018). Intriguingly, we detected that just before exiting the FP, dI1 growth cones very often sent a long protrusion into the FP (arrow, Fig. 5H, Movie 14). Live imaging clearly revealed that the activity and orientation of growth cones switched by about 90° after exiting the FP, as they flattened in the dorso-ventral axis and enlarged in the longitudinal axis *ex vivo* and *in vivo* (Fig. 5I,J, Movie 15). Another unexpected observation we made was that some dI1 growth cones transiently split while crossing the FP (asterisks in Movie 13). The splitting created two more or less equal branches (black arrows, Fig. 5K, Movie 16), but only one persisted and grew straight to the contralateral side, while the other one was retracted (black asterisks, Fig. 5K, Movie 16). Also this behavior was supported by snapshots from *in vivo* behavior of dI1 growth cones (arrows, Fig. 5L). Taken together, *ex vivo* live imaging combined with high magnification analysis of growth cone dynamics allowed us to characterize the behavior dI1 growth cones at choice points in more detail.

**Fig. 5.**
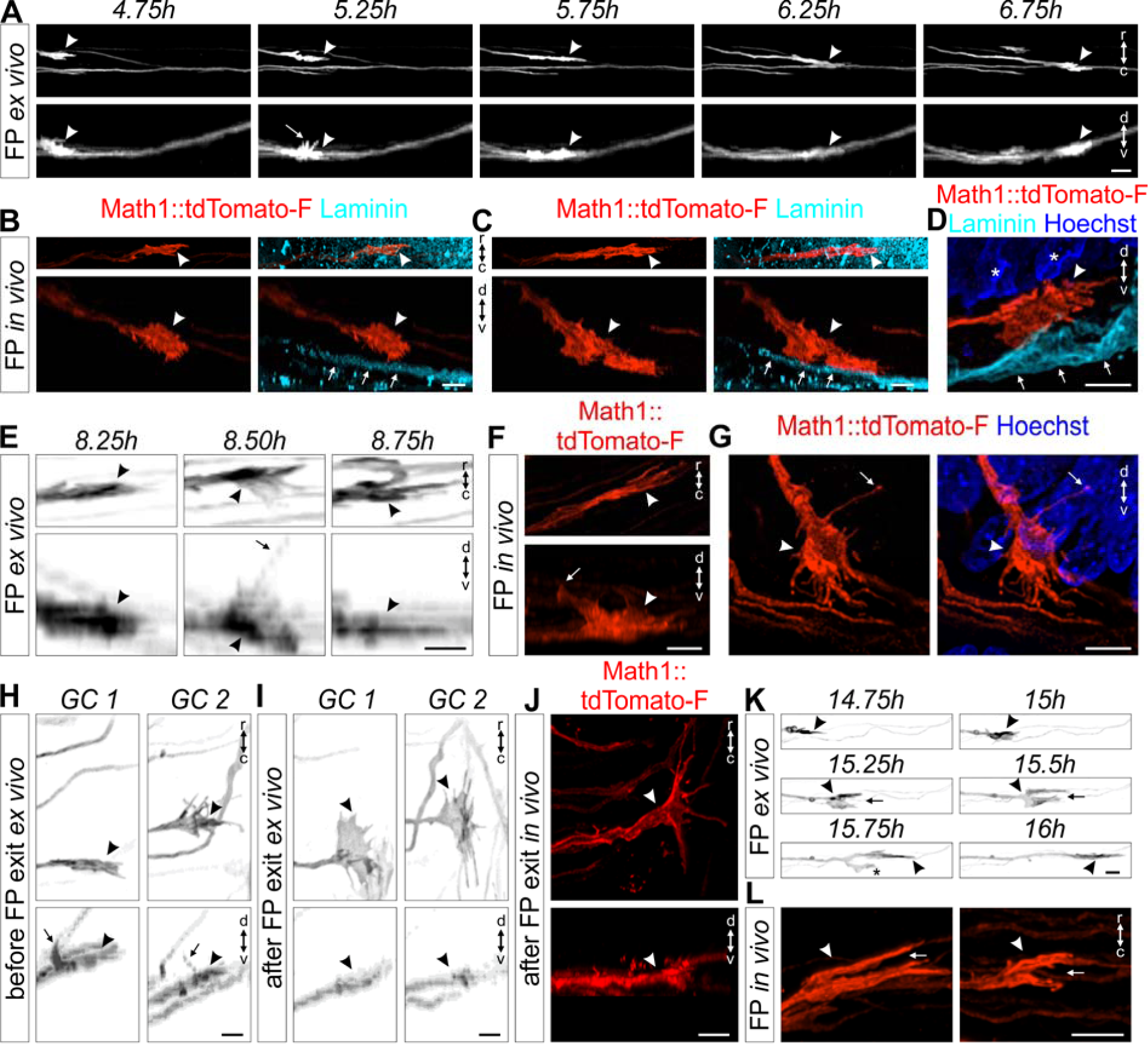
High magnification live imaging of dI1 commissural axons unraveled their orientation and activities at choice-points. (A) A Math1::tdTomato-F-positive growth cone (white arrowheads) crossing the FP shown in the rostro-caudal and dorso-ventral axis. White arrow shows the dorso-ventral orientation of the growth cone with some filopodia extended towards the apical FP. (B,C) Whole-mount immunostaining of Math1::tdTomato-F-positive dI1 growth cones in the FP at HH24.5 showed their dorso- ventral orientation *in vivo* (white arrowheads). The basal lamina was stained for laminin (white arrows). (D) A Math1::tdTomato-F-positive dI1 growth cone in the FP (white arrowhead) at HH22 showed its dorso-ventral orientation *in vivo* (white arrowheads). The basal lamina was stained for laminin (white arrows) and nuclei with Hoechst. White asterisk show nuclei from the FP cells. (E) Three consecutive snapshots from a time-lapse sequence showing the dorso-ventral activity of a Math1::tdTomato-F-positive growth cone crossing the FP (black arrowheads) with a long protrusion growing toward the FP soma level (black arrow). (F,G) A long protrusion growing towards the apical FP cell soma area (white arrows) could be also observed in dI1 growth cones crossing the FP *in vivo* (white arrowheads) after whole-mount staining of a HH24 spinal cord (F) or immunostaining on a HH22 spinal cord transverse section (G). (H) Example of two growth cones extracted from a time-lapse recording (black arrowheads) showing dorsal activity with a long protrusion (black arrows) growing towards the FP soma area just before exiting the FP. (I) The same growth cones shown in (H) after exiting the FP underwent a 90° change in their orientation. They now were thin in the dorso-ventral axis and enlarged in the rostro-caudal axis (black arrowheads). (J) HH24.5 whole-mount immunostaining of Math1::tdTomato-F-positive dI1 growth cone at the FP exit site showed that the orientation of post-crossing growth cones *in vivo* were the same as observed by live imaging (white arrowheads). (K) Example of a Math1::tdTomato-F- positive dI1 growth cone (black arrowhead) transiently splitting (black arrow) while crossing the FP. One branch always retracted (black asterisk) while the other one continued to grow straight to the contralateral side. (L) Split growth cones (white arrow) of Math1::tdTomato-F-positive dI1 axons (white arrowheads) could be observed *in vivo* after whole-mount staining of HH24.5 spinal cords. GC, growth cone; r, rostral; c, caudal; d, dorsal; v, ventral. Scale bars: 10 μm (A) and 5 μm (B-L).

### Live imaging unraveled the dynamics and morphologies of floor-plate cells during midline crossing

The orientation of dI1 growth cones as well as their behavior during FP crossing suggested that they have to squeeze their way between the basal feet of FP cells which are attached to the basal lamina (Yaginuma et al., 1991; Yoshioka and Tanaka, 1989). Moreover, very little was known about the morphology of FP cells during axonal midline crossing and their potential active contribution in this process has never been addressed (Campbell and Peterson, 1993; Yaginuma et al., 1991; Yoshioka and Tanaka, 1989). Therefore, we examined the behavior and morphology of FP cells during midline crossing in our *ex vivo* system. We electroporated spinal cords at HH17- 18 after injection of a plasmid encoding EGFP-F under the FP-specific Hoxa1 enhancer for expression of the membrane-bound fluorescent protein in FP cells (Li and Lufkin, 2000; Wilson and Stoeckli, 2011; Zisman et al., 2007) (Fig. 6A-D). With this we were able to see Hoxa1::EGFP-F-positive bulky FP basal feet in the commissure *in vivo* (white arrowheads, Fig. 6C) as well as their thin morphology and orientation (white arrows) that seemed to be tightly aligned with dI1 growth cones crossing the midline (white arrowheads, Fig. 6E). The morphology of medial FP basal feet with little extension in the rostro-caudal axis but enlarged in the dorso-ventral axis could be observed in real time using our *ex vivo* culture technique (Fig. 6F, Movie 17). Interestingly, we could observe dynamic protrusions sprouting from the basal feet in direction of axonal growth in the commissure (black arrows, Fig. 6F, Movie 17). Similar observations were made for basal feet of lateral FP cells (black arrow, Fig. 6G). They showed a very high activity with very dynamic protrusions towards the FP entry zone (black arrows, Fig. 6G, Movie 18). Importantly, we could observe a similar morphology of medial FP basal feet at the single-cell level *in vivo* (arrowheads, Fig. 6H, Movie 19). The dI1 commissural axons had the same orientation as the FP basal feet and seemed to grow in between these feet and interact with them *in vivo* (Movie 19 and 20). Moreover, we could observe similar protrusions coming either from lateral FP basal feet (white arrows) going towards pre-crossing dI1 axons arriving at the FP (white arrowheads, Fig. 6I), or from medial FP basal feet extending parallel to axons in the commissure (white arrows, Fig. 6J and Movie 21). The tight interaction between dI1 axons and FP basal feet during midline crossing was unexpected (Movie 21). The observation that protrusions from lateral FP basal feet (white arrowhead) were extending and contacting dI1 growth cones before they entered the FP (white arrows) suggested a much more active role of FP cells than anticipated (Fig. 6K, Movie 22). Last but not least, we could confirm that FP basal feet structures (yellow arrow) were present in between transiently splitting dI1 growth cones (white arrowheads and arrows) in the commissure (Fig. 6L, Movie 23). Taken together, these data showed for the very first time the detailed morphology of single FP cells, their dynamics and tight interactions with axons during midline crossing, and they emphasized the probable active role of FP cells in initiating contacts with growth cones before and during midline crossing.

**Fig. 6.**
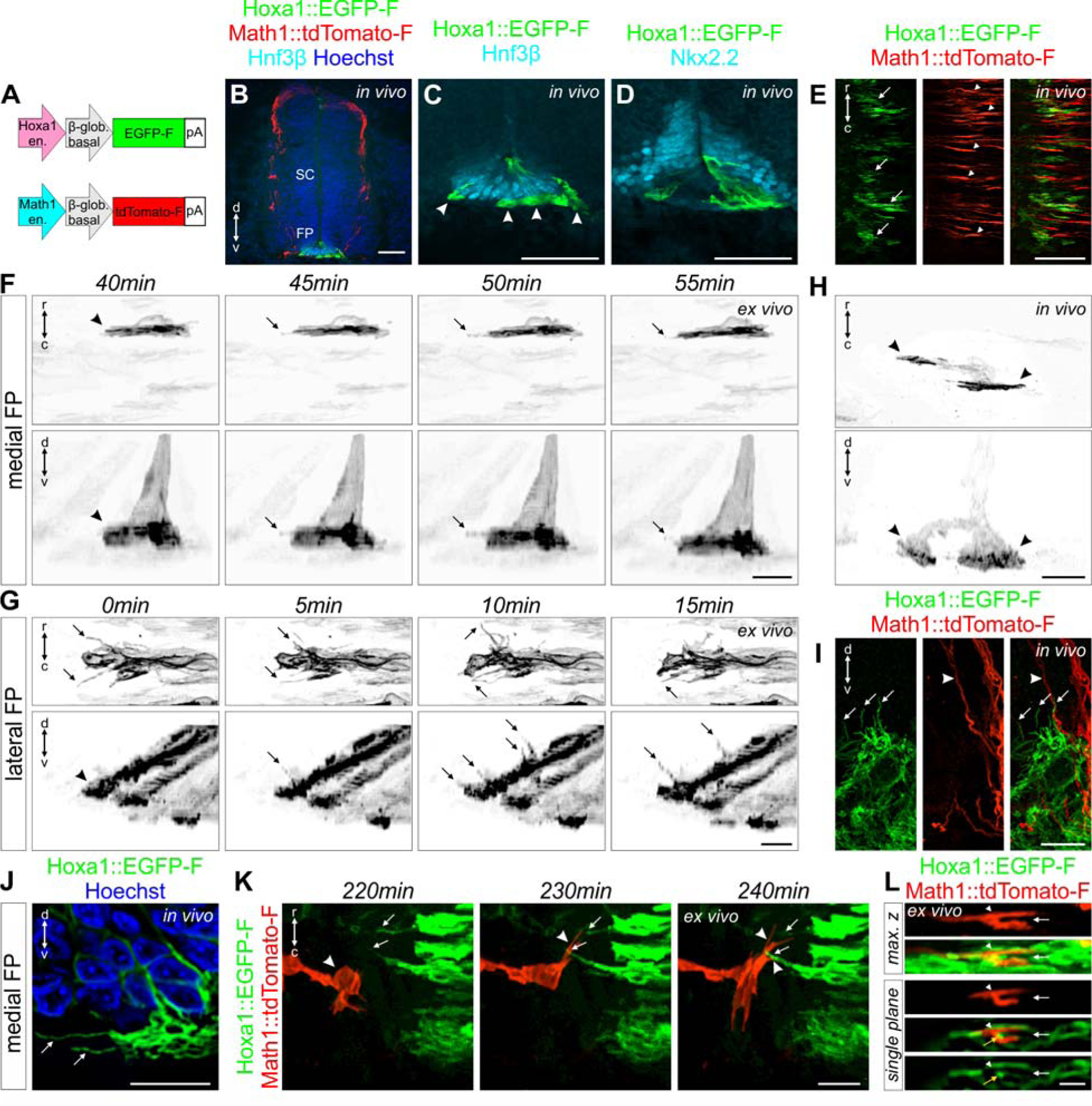
Live imaging of floor-plate cells during midline crossing shed light on their orientation and dynamics. (A) Schematic depicting plasmids that were electroporated at HH17-18 to visualize FP cells (Hoxa1 plasmid) together with dI1 axons (Math1 plasmid). (B) Immunostaining of HH22 spinal cord cryosections revealed the restricted expression of Hoxa1::EGFP-F in FP cells co-stained for Hnf3β. (C) Higher magnification of the section shown in (B) clearly showing Hnf3β-positive FP cells expressing EGFP-F. Bulky FP basal feet could be observed in the commissure (white arrowheads). (D) Immunostaining of HH22 spinal cord cryosections showing that EGFP-F expression driven by Hoxa1 enhancer was mostly absent in Nkx2.2-positive cells flanking the FP. (E) Whole-mount immunostaining of Math1::tdTomato-F-positive dI1 axons and Hoxa1::EGFP-F-positive FP cells revealed and alignment between dI1 growth cones (white arrowheads) and basal feet of FP cells (white arrows) in the commissure. (F) Example of a time-lapse recording of a single medial Hoxa1::EGFP-F-positive FP cell to reveal the geometry of its basal foot that was thin along the rostro-caudal axis but enlarged in the dorso-ventral axis (black arrowheads). The basal foot was highly dynamic with protrusions sprouting out in the directions of axonal growth in the commissure (black arrows). (G) Example of a time-lapse recording of lateral Hoxa1::EGFP-F-positive FP cells showing a very high activity of their basal feet with highly dynamic protrusions growing towards the arriving pre-crossing axons (black arrows). (H) Whole-mount immunostaining of a single medial Hoxa1::EGFP-F-positive FP cell *in vivo* at HH24.5 showing similar shape (black arrowheads) as observed by live imaging. Note that this FP cells contained two feet (black arrowheads). (I) Immunostaining of a HH22 spinal cord cryosection revealed that lateral Hoxa1::EGFP- F-positive FP cells also formed protrusions growing towards the pre-crossing axons *in vivo* (white arrows) where Math1::tdTomato-F-positive axons enter the FP (white arrowheads). (J) Similar protrusions were observed in HH22 medial FP basal feet in the commissure *in vivo* using immunostaining of cryosections (white arrows). The picture shows a single plane extracted from a Z-stack. (K) Snapshots extracted from a time- lapse sequence of a Math1::tdTomato-F-positive dI1 growth cone interacting (white arrowheads) with protrusions from Hoxa1::EGFP-F-positive FP basal feet (white arrows) before entering the FP. (L) Single snapshot from a time-lapse recording sequence showing maximum Z projection and single plane pictures of a transiently splitting (white arrow) Math1::tdTomato-F-positive dI1 growth cone (white arrowhead) in the FP with basal feet structures in between the two split branches (yellow arrowhead). SC, spinal cord; r, rostral; c, caudal; d, dorsal; v, ventral. Scale bars: 50 μm (A), 25 μm (B) and 10 μm (C-L).

### Trajectory and behavior of dI1 axons can be visualized in real time at choice points

Our *ex vivo* culture method not only offers great opportunities to characterize behavior of growth cones and FP cells in a preserved system, but it also opens new possibilities for tracking axonal behavior after specific perturbations of either the neurons or their environment. As example, the Wnt receptor Fzd3 (Frizzled-3) was specifically downregulated in Math1-positive dI1 neurons (Alther et al., 2016; Wilson and Stoeckli, 2011) (Fig. 7A). Fzd3 is required for the rostral turn of post-crossing commissural axons at the contralateral FP border *in vivo* (Alther et al., 2016; Lyuksyutova et al., 2003). We used our *ex vivo* culture system to visualize Math1::EGFP-F-positive dI1 axons expressing a microRNA for Fzd3 (miFzd3, Fig. 7B,C). This allowed us to follow in real time how dI1 growth cones were turning caudally instead of rostrally at the contralateral FP border (black arrowheads, Fig. 7B; Movie 24 and 25). Interestingly, many axons were found to turn erroneously in caudal direction at the same position, suggesting that axons were influenced by close contact with other axons (black asterisk and black arrows in Movie 24). In addition, some axons were found stalling at the FP exit site without initiating any turn (black arrowhead, Fig. 7C, Movie 25). Nonetheless, the growth cones remained highly dynamic. The changes in morphology were accompanied with transient retraction and re-extension but without a clear change in directionality. Importantly, expressing a control microRNA (mi2Luc) did not impact the guidance of dI1 axons at the contralateral FP border (Fig. 7D, Movie 26). Taken together, our method can be used to study the behavior of axons after perturbation of candidate genes specifically in the neurons or their target in real time.

**Fig. 7.**
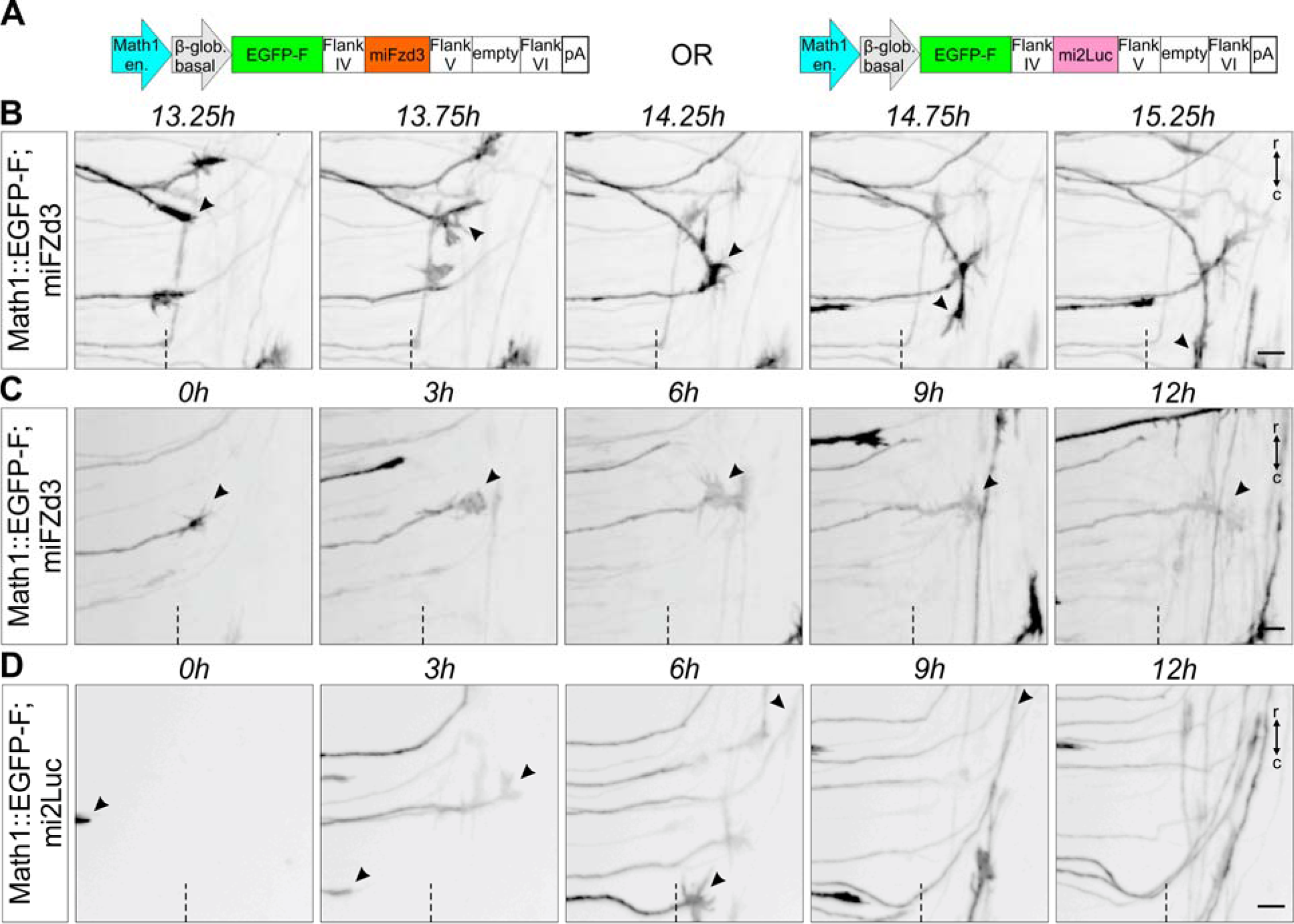
Live imaging after dI1 neuron-specific knockdown can be used to visualize mutant axons in intact spinal cord preparations. (A) Schematics depicting the plasmid constructs used to knockdown Fzd3 in dI1 neurons. A plasmid expressing a microRNA against Luciferase (mi2Luc) was used as a control. (B) Time-lapse sequence showing a dI1 commissural axon turning caudally instead of rostrally at the FP contralateral border after silencing Fzd3 (black arrowheads). (C) Time-lapse sequence showing a dI1 commissural axon stalling at the contralateral FP border after silencing Fzd3 (black arrowheads). The growth cone kept remodeling but was not able to turn in either direction. (D) Time-lapse sequence showing dI1 axons expressing a microRNA against luciferase. These axons were not impacted and after exiting the FP they all turned rostrally (black arrowheads). Dashed black line represents the FP exit site. r, rostral; c, caudal. Scale bars: 10 μm.

## DISCUSSION

The possibility to follow one axon over time, rather than deducing behavior from snapshots of different axons, allowed us to extract detailed information on the timing of midline crossing and the tight interaction between FP cells and the growth cones in a higher vertebrate model (Figure 8). Our *ex vivo* system offers significant improvement over existing open-book culture systems which lead to obvious guidance artefacts at the contralateral FP border (Figure 2) (Pignata et al., 2019). Our *ex vivo* culture system is highly reproducible and generates a manageable amount of data compared to live imaging using light sheet-based microscopy, for example (Liu et al., 2018).

**Fig. 8.**
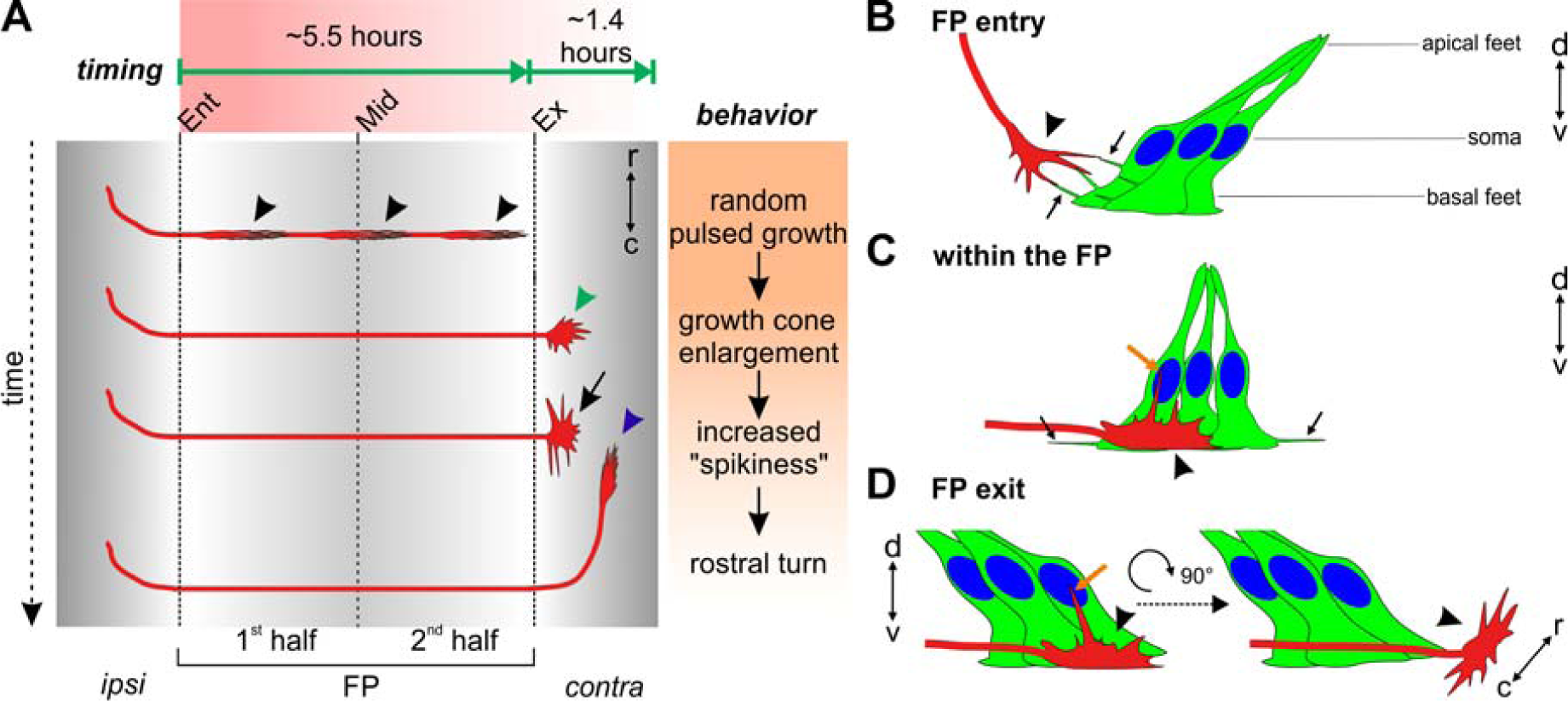
Cartoon depicting the midline crossing characteristics of dI1 axons based on data extracted from our *ex vivo* culture system. (A) On average, it took dI1 axons 5.6 hours to cross the midline. Growth cones showed a random pulsed growth and had a thin shape in the growth direction (black arrowheads). At the FP exit site, dI1 growth cones were first enlarged (green arrowhead), then extended filopodia along the longitudinal axis (black arrow) right before turning rostrally (blue arrowhead). After arriving at the exit site of the FP, it took dI1 axons on average about 1.4 hours to turn rostrally. In fact, the first exiting dI1 axons took longer to turn rostrally than the late exiting ones. (B-D) Live imaging of intact spinal cords ex ovo using a high magnification objective shed light on dI1 growth cone orientation, FP morphology and dynamics during midline crossing. (B) While dI1 growth cones (black arrowhead) approached the FP, basal feet of lateral FP cells sent protrusions towards them and eventually interacted with them (black arrows). (C) When dI1 growth cones crossed the FP (black arrowhead), their dorso-ventral orientation aligned perfectly with the orientation of basal feet of medial FP cells. While basal feet of FP cells sent protrusions in axonal growth direction (black arrows), dI1 growth cones sent long filopodia in direction of the apical FP, towards the FP cell soma (orange arrow). (D) Just before exiting the FP, dI1 growth cones showed dorso-ventral activity with a long protrusion growing towards the FP soma (orange arrow) area followed by a 90° change in their orientation to become flattened in the dorso-ventral axis and enlarged in the longitudinal axis (black arrowhead). Ent, entry; Mid, midline; Ex, exit; r, rostral; c, caudal; ipsi, ipsilateral; contra, contralateral.

Our comparative analyses demonstrate that midline crossing of dI1 axons in our *ex vivo* system was very similar to what happens *in vivo* (Figure 2,4,5 and S4). Therefore, our *ex vivo* system can be used to monitor and assess axonal behavior at choice points. We could detect that dI1 growth cones took on average 5.6 hours to cross the entire FP and that they did so in a pulsed manner (black arrowheads, Figure 8). We could also measure that they needed on average 1.4 hours to initiate their rostral growth, and that the first axons exiting the FP took longer than the followers (Figure 8A). In total they needed almost 7 hours from entering the FP to making the decision to turn rostrally (blue arrowhead, Figure 8A). This is enough time for growth cones to change their responsiveness to specific guidance cues for crossing and exiting the FP as well as for turning rostrally due to changes in receptor expression regulated at the post-translational, translational and even transcriptional level (Nawabi et al., 2010; Philipp et al., 2012; Pignata et al., 2019; Preitner et al., 2016; Stoeckli, 2018; Wilson and Stoeckli, 2013).

The growth cone is the decision center where axon guidance instructions are transduced to the cytoskeleton (Vitriol and Zheng, 2012). With our newly developed *ex vivo* system, dynamic changes in dI1 commissural growth cone morphology and behavior at the midline can be observed in real time. Growth cones were thin and elongated in the FP with their major extension in the dorso-ventral axis (black arrowheads, Figure 8A,C). At the FP exit site, they showed a 90° rotation to be enlarged and active in the longitudinal axis (green arrowhead and black arrow, Figure 8A, black arrowhead, Figure 8D). The fact that dI1 growth cones sent a long filopodium into the FP, towards the FP cell soma area, while crossing it and just before exiting it, suggests that they might need to read signals from this area in order to move on and exit the FP (orange arrows in Figure 8C and D). The extension of long filopodia just before FP exit and rostral turning suggests that actin polymerization might be required to sense repulsive cues – for instance SlitN and Shh, respectively – and transduce the signal into the growth cone, as suggested for Slit-induced growth cone collapse *in vitro* (McConnell et al., 2016). Further investigations using our *ex vivo* culture system will be required to understand the role of cytoskeletal dynamics in axonal navigation of the intermediate target.

Our method also suggested a probable active contribution of FP cells to axon guidance, as we found the cells of the intermediate target to by very dynamic and to extend protrusions in directions of the arriving axons, or to actively engage with axons in the FP. Thus, it seems that the intermediate target is much more than a passive by-stander and provider of attractive and repulsive axon guidance and cell adhesion molecules. We characterized the FP cell morphologies in detail in the medial as well as the lateral FP. Basal feet appeared to be enlarged and oriented parallel to commissural growth cones (Figure 8C). The lateral FP basal feet sent protrusions (black arrows) towards dI1 growth cones approaching the FP and eventually interacted with them (black arrowhead, Figure 8B). This intriguing observation led us to speculate whether these protrusions might be cytonemes. Cytonemes are long protrusions known to spread and deliver morphogens, such as Wnts and Shh, to neighboring or more distant cells (González-Méndez et al., 2019; Sanders et al., 2013; Stanganello and Scholpp, 2016). Given the fact that Shh is involved in guiding pre-crossing commissural axons towards the FP and that Shh and Wnts are both involved in guiding post-crossing axons towards the brain at the contralateral FP border, it is tempting to speculate that these protrusions might deliver such signals to the growth cones at choice points (Avilés et al., 2013). Moreover, we could appreciate how much the axons and their growth cones were intermingled within the medial FP basal feet which also formed long dynamic protrusions within the commissure (black arrowheads, Figure 8C). In sum, the combination of our live imaging approach with a FP-specific marker will give the opportunity to further characterize the behavior of intermediate target cells with regard to axon guidance at choice points.

Ultimately, our method will be useful to get more insights into molecular mechanisms of axon guidance at a choice point, when combined with *in ovo* RNAi for specific gene knockdowns either in the neurons or in their environment, as exemplified with Fzd3 knockdown experiments (Figure 7) (Andermatt et al., 2014; Pekarik et al., 2003). Similarly, pharmacological blockers will permit to screen for components required downstream of growth cone receptors to transduce guidance signals. Usually such experiments are conducted *in vitro* with cultured neurons growing axons in a very artificial environment. Thus, our method offers the advantages of an *in vitro* experiment in an intact complex ‘*in vivo*-like’ environment. The use of specific reporters will also allow for the assessment of dynamic changes of second messengers or the actin cytoskeleton in growth cones for example (Nicol et al., 2011; Nichols and Smith, 2019). Moreover, the use of other sets of enhancers and promoters might offer the possibility to study the dynamics of midline crossing in other subtypes of commissural neurons in the spinal cord and in the brain (Hadas et al., 2014; Kohl et al., 2012).

## MATERIALS AND METHODS

### *In ovo* electroporation

Plasmids encoding farnesylated td-Tomato under control of the Math1 enhancer and the β-globin promoter for dI1 neuron-specific expression (Math1::tdTomato-F, 700 ng/μl) and farnesylated EGFP under control of the β-actin promoter (β-actin::EGFP-F, 30 ng/μl) were co-injected into the central canal of the chicken neural tube *in ovo* at HH17-18 (Hamburger and Hamilton, 1951) and unilaterally electroporated, using a BTX ECM830 square-wave electroporator (five pulses at 25 V with 50 ms duration each), as previously described (Wilson and Stoeckli, 2012) (Fig. 1A,B). A final concentration of 0.1% (vol/vol) of Fast Green was added to the plasmid mix to trace injection site and volume of the plasmid mix. After electroporation, embryos were covered with sterile PBS and eggs were sealed with tape and incubated at 39°C for 26-30 hours, until embryos reached stage HH22, or for 36-46 hours, until embryos reached HH24. For the FP study EGFP-F was expressed from a plasmid with the Hoxa1 enhancer and the β- globin minimal promoter (Wilson and Stoeckli, 2011) (Hoxa1::EGFP-F, 1000 ng/μl) and co-injected with the Math1::tdTomato-F plasmid (700 ng/μ electroporated as above, or bilaterally electroporated (3 pulses in each direction at 25 V with 50 ms duration each). For knockdown of Fzd3 (or luciferase as control) in dI1 neurons (Math1 enhancer) plasmids previously published (Math1::EGFP-F; miFzd3 and μ -actin::mRFP plasmidl) and unilaterally electroporated at HH16 for more efficient knockdown (Alther et al., 2016).

### Dissection of intact spinal cords

Intact spinal cords were dissected from HH22 embryos in ice-cold, sterile PBS (Gibco) in a silicon-coated Petri dish with sterile instruments. Embryos were pinned down with their dorsal side down with thin needles (insect pins). Here, special care was taken not to damage or detach meninges surrounding the spinal cord by avoiding too much rostro-caudal and lateral tension. Internal organs and ventral vertebrae were removed to access the spinal cord. Ventral roots were cut off and the spinal cord was carefully extracted from the embryo with forceps, avoiding any excessive bending. Note that dorsal root ganglia were not cut off and all were still attached to the spinal cord. The ventral and dorsal midline were kept intact throughout dissection. Finally, remaining dorsal tissues were discarded. See Fig. S1 for a detailed step-by-step protocol to successfully dissect intact HH22 spinal cords. Note that this procedure can also be applied to older embryos (at least HH24-25). Once intact spinal cords were dissected and cleaned from any remaining dorsal tissues they were embedded with the ventral side down in a warm (39°C) 100-µl drop of 0.5% low-melting agarose (FMC, Fig. 1D,E, Fig. S1G) containing a 6:7 ratio of spinal cord medium [MEM with Glutamax (Gibco) supplemented with 4 mg/ml Albumax (Gibco), 1 mM pyruvate (Sigma), 100 Units/ml Penicillin and 100 µg/ml Streptomycin (Gibco)] in a 35-mm Ibidi µ-Dish with glass bottom (Ibidi, #81158). Note that the spinal cord should be as straight as possible with the dorso-ventral axis perpendicular to the glass bottom, as any pronounced curvature or tilting of the midline would induce axon guidance artefacts or death of the axons, respectively. Once the agarose solidified (around 5 min at room temperature), 200 µl of spinal cord medium were added to the drop and the culture could be started. A 12-mm flexiPERM conA ring (Sarstedt) was placed in the center of the culture dish before the agarose drop was added (the drop should not touch the ring). Hence, the medium added to the drop of low-melting agarose could touch the ring all around and therefore stabilize the position of the agarose drop and avoid any movement of the spinal cord during recordings thanks to surface tension (Fig. S1G).

### Dissection of open-books

Open-book preparations of spinal cords were dissected from HH24 embryos as previously described in a video protocol for HH25-26 embryos (Wilson and Stoeckli, 2012). The first steps were identical to the protocol for intact spinal cord dissection given above (steps A,B in Fig. S1). Starting there, the tension along the rostro-caudal axis was increased using the upper and lower needles and meninges were removed with a blade made of fire-polished tungsten wire. Spinal cords were cut transversally at the wing and leg levels and carefully extracted from the embryo with forceps. At this point, the dorsal midline spontaneously opened. Open-book preparations of spinal cords were then plated with the apical side down (Fig. 1G,H) in the center of a 35-mm Ibidi µ- Dish with glass bottom (Ibidi, #81158), pre-coated with 20 µg/ml poly-L-lysine (Sigma). A homemade, harp-like holder made out of a Teflon ring and thin nylon strings was used to keep the spinal cord in place (Fig. 1G). Note that the strings were barely touching the open-books but stabilized the flat position of the spinal cord. Then, 100 µl of 0.5% low-melting agarose (FMC) containing a 6:7 ratio of spinal cord medium were added on top of the spinal cord. Once the agarose solidified (around 5 min at room temperature), 200 µl of spinal cord medium were added to the agarose drop and the culture could be started.

### Live imaging

Live imaging recordings were carried out with an Olympus IX83 inverted microscope equipped with a spinning disk unit (CSU-X1 10,000 rpm, Yokogawa). Cultured spinal cords were kept at 37°C with 5% CO_2_ and 95% air in a PeCon cell vivo chamber (PeCon). Temperature and CO_2_-levels were controlled by the cell vivo temperature controller and the CO_2_ controller units (PeCon). Spinal cords were incubated for at least 30 min before imaging was started. We acquired 18-40 planes (1.5 µm spacing) of 2x2 binned z-stack images every 15 min for 24 hours with a 20x air objective (UPLSAPO 20x/0.75, Olympus) and an Orca-Flash 4.0 camera (Hamamatsu) with the Olympus CellSens Dimension 2.2 software. We performed most of our recordings in the lumbar level of the spinal cord and always took 3 channels of interest: emission at 488 nm and 561 nm, as well as brightfield. Recordings of Fzd3 or luciferase knockdown axons were performed at the thoracic level. For higher magnification recordings, a 40x silicone oil objective was used (UPLSAPO S 40x/1.25, Olympus) with same acquisition settings as above and images taken every 5-15 min.

### Data processing and virtual tracing

Z-stacks and maximum projections of Z-stack movies were evaluated and processed using Fiji/ImageJ (Schindelin et al., 2012). The MtrackJ plugin (Meijering et al., 2012) was used to virtually trace single Math1-positive dI1 commissural axons crossing the FP. This helped to keep track of which axons had already been quantified. The leading edge (and not filopodia) of growth cones was always selected for each time point. At the exit site, as growth cones very often slightly changed their directionality and drastically change their shape before turning, the central domain of the growth cone was selected with the tracing tool. Only axons that entered, crossed and exited the FP during the 24- hour imaging period were traced and quantified. Overlays of labeled axons with EGFP-F and brightfield channels were used to assess the FP boundaries and midline localization. The virtual tracing tool was also used to extract the instantaneous growth speed for each single axon. Note that the montage of dI1 commissural axons shown in Movie 2 was generated from z-stacks that were 2D deconvolved (nearest neighbor) using the Olympus CellSens Dimension 2.2 software and assembled with Fiji/ImageJ. All data acquired with higher magnification (40x silicone oil objective) were 3D deconvolved using constrained iterative deconvolution of the Olympus CellSens Dimension 2.2 software (5 iterations with adaptive PSF and background removal, Olympus). Maximum projections of live images containing Hoxa1::EGFP-F-positive cells (channel) were corrected for photo bleaching in Fiji/ImageJ.

### Temporal-color projections and kymographic analysis

Temporal-color projections were generated using Fiji/ImageJ. Kymograph analysis of axons crossing or exiting the FP as previously described (Medioni et al., 2015) using a region of interest (ROI) selection, the re-slice function and the z-projection of the re- sliced results in Fiji/ImageJ, which allowed following pixel movements within the horizontal axis. The ROI in the FP was selected as a 103x51 µm^2^ (Fig. 3C,D) or 103x27 µm^2^ (Fig. 3E,F) rectangle.

### Immunohistochemistry

Spinal cords dissected from HH22 embryos or intact spinal cords that were cultured for one day *ex vivo* were fixed one hour at room temperature with 4% paraformaldehyde in PBS, washed 3 times for 5 min each with PBS and cryopreserved for at least 24h at 4°C in 25% sucrose in PBS. After mounting in O.C.T. compound (Tissue-Tek) and freezing the spinal cords, 25-µm thick cryosections were collected using a cryostat. The next day, sections were blocked and permeabilized 1h at room temperature with 5% FCS in 0.1% Triton X-100 in PBS (blocking buffer). Primary antibodies were diluted in blocking buffer and added to sections overnight at 4°C (1:400 for goat-anti-GFP-FITC, Rockland; 1:2,500 for rabbit-anti-RFP, antibodies-online; supernatants of monoclonal antibodies obtained from DSHB: anti-Lhx2 (clone 1C11), anti-islet-1, (clone 40.2D6), anti-Nkx2.2 (clone 74.5A5), anti-Hnf3β (clone 4C7); 3.1 µg/ml of mouse-anti-Shh, clone 5E1). The next day, sections were washed 3 times for 15 min each at room temperature with 0.1% Triton-X100 in PBS. Primary antibodies (except anti-GFP-FITC) were detected with a 2- hour incubation in adequate secondary antibodies diluted in blocking buffer (1:1,000 for donkey-anti-mouse-IgG-Cy5 or donkey-anti-rabbit-IgG-Cy3, Jackson ImmunoResearch). Finally, nuclei were counterstained for 10 min at room temperature with 2.5 µg/ml of Hoechst diluted in 0.1 Triton X-100 in PBS (Invitrogen), washed 3 times for 10 min each with 0.1% Triton X-100 in PBS and 2 times for 5 min each with PBS before mounting the slides in Mowiol/DABCO. Images were taken with an Olympus IX83 inverted microscope equipped with a spinning disk unit (CSU-X1 10,000 rpm, Yokogawa), a 20x air objective (UPLSAPO 20x/0.8, Olympus) or a 40x silicon oil objective (UPLSAPO S 40x/1.25, Olympus), and an Orca-Flash 4.0 camera (Hamamatsu) with the Olympus CellSens Dimension 2.2 software, or with an Olympus BX61 upright microscope and a 10x air objective (UPLFL PH 10x/0.30, Olympus) or 40x water objective (UAPO W/340 40x/1.15, Olympus) and an Orca-R^2^ camera (Hamamatsu) with the Olympus CellSens Dimension 2.2 software.

### Whole-mount immunostaining

Intact spinal cords dissected from HH24-25 embryos were fixed for 1h at room temperature in 4% paraformaldehyde in PBS and washed 3 times 10 min each with PBS. Spinal cords were permeabilized and incubated for 1h at room temperature with 5% FCS in 0.1% Triton X-100 in PBS (blocking buffer). Primary antibodies were diluted in blocking buffer and added to spinal cords for incubation overnight a 4°C (1:800 of goat-anti-GFP-FITC, Rockland; 1:5,000 of rabbit-anti-RFP, antibodies-online). The next day, sections were washed 3 times 30 min each at room temperature with 0.1% Triton- X100 in PBS. The primary antibody against RFP was detected with a 2-hour incubation in diluted donkey-anti-rabbit-IgG-Cy3 antibody in blocking buffer (1:1,000, Jackson ImmunoResearch). Finally, nuclei were counterstained for 30 min at room temperature with 2.5 µg/ml of Hoechst diluted in 0.1 Triton X-100 in PBS (Invitrogen). Samples were washed 3 times 30 min each with 0.1% Triton X-100 in PBS and 2 times 15 min each in PBS. Stained spinal cords where mounted in 100 µl of 0.5% low-melting agarose in PBS with the ventral midline pointing down on a 35-mm Ibidi µ-Dish with glass bottom (Ibidi, #81158), similarly as described above. This allowed accessing the commissure of fixed *in vivo* samples and visualization of dI1 axons and their growth cone with an Olympus IX83 inverted microscope equipped with a spinning disk unit (CSU-X1 10,000 rpm, Yokogawa). Pictures were taken with a 4x air objective (UPLFLN PH 4x/0.13, Olympus), a 20x air objective (UPLSAPO 20x/0.75, Olympus)) or a 40x silicon oil objective (UPLSAPO S 40x/1.25, Olympus) and an Orca-Flash 4.0 camera (Hamamatsu) with the Olympus CellSens Dimension 2.2 software. Note that the same mounting and microscopy procedure was applied to HH23-25 intact spinal cords that were not stained and were used for the quantification of average growth cone areas *in vivo* (shown in Fig. 4C).

### Statistics and Figures assembly

Statistical analyses were carried out with GraphPad Prism 7.02 software. All data were assessed for normality (normal distribution) using the D’Agostino and Pearson omnibus K2 normality test and visual assessment of the normal quantile-quantile plot before choosing an appropriate (parametric or non-parametric) statistical test. P values of the simple linear regression shown in Fig. 3I,J demonstrate whether the slope is significantly different to zero and the dashed lines represent the 95% confidence intervals. Figures were assembled using Corel Draw 2017.

## Acknowledgements

We thank Dr. Beat Kunz for excellent technical assistance.

## Competing interests

The authors declare no competing interests.

## Author contributions

AD, designed experiments, performed experiments, evaluated data and contributed to the writing of the manuscript, NRZ, generated the Hoxa1 plasmid, ETS, designed experiments, supervised the project and contributed to the writing of the manuscript.

## Funding

This project was supported by the Swiss National Science Foundation.

---

**Movie 1. 24-hour time-lapse recording of the *ex vivo* cultured intact spinal cord shown in** Fig. 2A. Math1::tdTomato-F-positive dI1 axons (red), β positive axons and cells (green) and brightfield maximum projections are shown in parallel. One z-stack was taken every 15 minutes for each channel. White and orange dashed lines indicate FP boundaries and the midline, respectively. Rostral is up.

**Movie 2. 8-hour segment of a time-lapse recording of an intact spinal cord cultured *ex vivo*.** Math1::tdTomato-F-positive dI1 axons are shown in black. Maximum projections of z-stacks taken every 15 minutes are represented as well as 3D rotations at different time points (at FP entry, during FP crossing, at FP exit and after rostral turn). Black and orange dashed lines indicate FP boundaries and midline, respectively. Rostral is up.

**Movie 3. 4-hour time-lapse recording of the *ex vivo* cultured open-book preparation of the spinal cord shown in Fig. 2C,D**. Math1::tdTomato-F-positive dI1 -actin::EGFP-F-positive axons and cells (green) and brightfield maximum β projections are shown in parallel. One z-stack was taken every 15 minutes for each channel. White and orange dashed lines indicate FP boundaries and the midline, respectively. Rostral is up.

**Movie 4. 24-hour time-lapse recording of an *ex vivo* cultured intact spinal cord used for kymographic analysis in** Fig. 3D. Math1::tdTomato-F-positive dI1 axons are shown in black. Maximum projections of z-stacks taken every 15 minutes are represented. Black and orange dashed lines indicate FP boundaries and midline, respectively. Rostral is up.

**Movie 5. 7.75-h time-lapse recording of the segment shown in** Fig. 3G. This movie shows the virtual tracing of the growth cone (black arrowhead) at each time point revealing the instantaneous growth speed. Math1::tdTomato-F-positive dI1 axons are shown in black. Maximum projections of z-stacks taken every 15 minutes are represented. Black and orange dashed lines indicate FP boundaries and midline, respectively. Rostral is up.

**Movie 6. Example of growth cone enlargement at the floor-plate exit site.** Growth cone of a Math1::tdTomato-F-positive dI1 commissural axon (black arrowheads) exiting the FP (black dashed line) and becoming spiky with many filopodia before turning rostrally (black arrow). Rostral is up.

**Movie 7. Example of growth cone enlargement at the floor-plate exit site.** Two growth cones of Math1::tdTomato-F-positive dI1 commissural axons (black arrowheads) exiting the FP (black dashed line) and becoming spiky with filopodia growing in rostral and caudal direction before turning rostrally (black arrows). Rostral is up.

**Movie 8. Example of growth cone enlargement and transient splitting at the floor- plate exit site.** Growth cone of a Math1::tdTomato-F-positive dI1 commissural axon (black arrowheads) exiting the FP (black dashed line), becoming spiky with a lot of filopodia and transiently splitting before turning rostrally (black arrow). Rostral is up.

**Movie 9. 24-hour time-lapse recording with high magnification objective of cultured intact spinal cord.** Math1::tdTomato-F-positive dI1 axons are shown in black. Maximum projections of z-stacks taken every 15 minutes are represented. Dashed lines represent FP boundaries. Rostral is up.

**Movie 10. Time-lapse recording within the floor plate uncovered dI1 growth cones orientation.** Math1::tdTomato-F-positive dI1 growth cone (white arrowhead) crossing the FP showed a dorso-ventral orientation with filopodia activities in the dorso-ventral axis (white arrows). Maximum projections of z-stacks (rostro-caudal and dorso-ventral axis) taken every 15 minutes are represented as well as a 3D rotation. r, rostral; c, caudal; d, dorsal; v, ventral.

**Movie 11. dI1 growth cones showed the same dorso-ventral orientation in the floor plate *in vivo* (whole-mount staining of HH24.5 intact spinal cord).** 3D representation of two Math1::tdTomato-F-positive dI1 growth cones (white arrowheads) showing little extension in the rostro-caudal axis but enlarged size in the dorso-ventral axis. The basal lamina was stained for laminin (shown in cyan, white arrows). r, rostral; c, caudal; d, dorsal; v, ventral.

**Movie 12. Early crossing dI1 growth cones showed the same dorso-ventral orientation in the floor plate *in vivo* as observed *ex vivo* (immunostaining of HH22 spinal cord cross-sections).** 3D representation of a Math1::tdTomato-F-positive dI1 growth cones (white arrowheads) showing little extension in the rostro-caudal axis but enlarged size in the dorso-ventral axis. The basal lamina was stained for laminin (shown in cyan) and nuclei counter-stained with Hoechst (shown in blue). r, rostral; c, caudal; d, dorsal; v, ventral.

**Movie 13. 14-hour time-lapse recording of dI1 axons crossing the floor-plate.** Rostro-caudal, dorso-ventral and 3D representations of Math1::tdTomato-F-positive dI1 axons crossing the FP. dI1 growth cones transiently split in the FP (black asterisks) and showed filopodial extensions towards the FP cell soma area (black arrowheads). r, rostral; c, caudal; d, dorsal; v, ventral.

**Movie 14. Detailed analysis of dI1 growth cone shape at the floor-plate exit site in real time.** Sequence from a time-lapse recording showing two Math1::tdTomato-F- positive dI1 commissural growth cones exiting the FP (black arrowheads). Both sent a protrusion towards the FP cell soma just before exiting (black arrows) and changed their orientation upon exiting the FP by 90° to adopt an enlarged size in the longitudinal axis. Dashed line represents the FP exit. r, rostral; c, caudal; d, dorsal; v, ventral.

**Movie 15. dI1 growth cone at the floor-plate exit site of a HH24.5 spinal cord *in vivo*.** Example of a Math1::tdTomato-F-positive growth cone (white arrowhead) with little extension along the dorso-ventral axis but enlarged in the rostro-caudal axis. r, rostral; c, caudal; d, dorsal; v, ventral.

**Movie 16. dI1 growth cone transiently split in the floor plate.** Sequence from a time-lapse recording showing a Math1::tdTomato-F-positive dI1 commissural growth cone crossing the FP (black arrowhead) and transiently splitting (black arrowhead). Note that one branch retracted (black asterisk) while the other continued to grow straight towards the contralateral side (black arrowhead). Rostral is up.

**Movie 17. Time-lapse recording sequence of a Hoxa1::EGFP-F-positive medial floor plate cell.** Live imaging of a medial FP cell showed that it had a thin basal foot in the rostro-caudal axis. In the dorso-ventral axis to foot was large. The FP cell extended highly dynamic protrusions from its foot in both directions of axonal growth (black arrows). r, rostral; c, caudal; d, dorsal; v, ventral.

**Movie 18. Time-lapse recording sequence of Hoxa1::EGFP-F-positive lateral floor- plate cells.** Live imaging of lateral FP cells revealed a very high activity of their basal feet with highly dynamic protrusions sprouting out in the direction of the arriving pre-crossing axons (black arrows). r, rostral; c, caudal; d, dorsal; v, ventral.

**Movie 19. Whole-mount staining of a medial floor-plate cell uncovered its shape *in vivo*.** Example of a single Hoxa1::EGFP-F-positive medial FP cell at HH24.5 (shown in green) with two thin basal feet in the rostro-caudal axis that were enlarged in the dorso-ventral axis at the level of the commissure (white arrows). The orientation of basal feet was aligned to a Math1::tdTomato-F-positive dI1 growth cone (white arrow, shown in red). r, rostral; c, caudal; d, dorsal; v, ventral.

**Movie 20. Whole-mount staining of medial floor-plate cells revealed alignment between their basal feet and dI1 growth cones *in vivo*.** Example of Hoxa1::EGFP-F- positive medial FP cells at HH24.5 (shown in green) with basal feet oriented in the dorso-ventral axis (white arrows). Math1::tdTomato-F-positive dI1 growth cones were aligned with the FP basal feet (white arrow, shown in red). r, rostral; c, caudal; d, dorsal; v, ventral.

**Movie 21. Z-stack animation of the medial floor-plate area reveal protrusions of basal feet and demonstrated close interaction between dI1 axons and floor-plate basal feet in the commissure *in vivo*.** Immunohistochemistry of HH22 spinal cord sections revealed that Hoxa1::EGFP-F-positive medial FP basal feet (shown in green) also contained protrusions in the commissure *in vivo* (white arrows). FP basal feet seemed to enwrap Math1::tdTomato-F-positive axons in the commissure (white arrowheads). Nuclei were counterstained with Hoechst (shown in blue). d, dorsal; v, ventral.

**Movie 22. Time-lapse recording sequence of a dI1 commissural growth cone interacting with protrusions from floor-plate basal feet before entering the floor plate.** A Math1::tdTomato-F-positive dI1 growth cone (shown in red) extended filopodia in the direction of the FP (white arrowheads) and eventually interacted with protrusions (white arrows) coming from the Hoxa1::EGFP-F-positive FP basal feet (shown in green) before entering the FP. Note that at “t=240 min” a Z-stack animation is shown to pinpoint the close vicinity between basal feet protrusions and the growth cone (white arrows and arrowheads, respectively). Rostral is up.

**Movie 23. Time-lapse recording sequence of a dI1 commissural growth cone transiently splitting in the commissure in between floor-plate basal feet.** A Math1::tdTomato-F-positive dI1 growth cone (shown in red, white arrowhead) growing through the FP basal feet (shown in green) and undergoing transient splitting (white arrow). Note that at “t=2 h” (splitting time point) a Z-stack animation is shown to pinpoint that FP basal feet structures (yellow arrow) were present in between both branches of the split dI1 growth cone. Rostral is up.

**Movie 24. 48-hour time-lapse recording sequence showing aberrant caudal turning of dI1 commissural axons after silencing Fzd3.** Math1::EGFP-F; miFzd3- positive dI1 axons exiting the FP are shown in black. The turning behavior at the contralateral FP border was randomized with a substantial number of mutant dI1 axons turning caudally instead of rostrally (black arrows). Often, collective behavior was seen, that is, after a first axon turning caudally many other growth cones seemed to follow the same path and fasciculated with axons that turned in the wrong direction (black asterisk). Dashed line represents the FP exit site. Rostral is up.

**Movie 25. 48-hour time-lapse recording showing growth cone stalling at the floor- plate exit site after silencing Fzd3.** Math1::EGFP-F; miFzd3-positive dI1 axons exiting the FP are shown in black. Black arrowhead shows a mutant dI1 axons stalling at the contralateral FP border. Some axons were also turning caudally (black arrows). Dashed line represents the FP exit site. Rostral is up.

**Movie 26. 48-hour time-lapse recording showing normal behavior of control- treated dI1 axons at floor-plate exit.** Math1::EGFP-F; mi2Luc-positive dI1 axons exiting the FP are shown in black. All growth cones behaved normally at the contralateral FP border and turned rostrally (black arrowheads). Dashed line represents the FP exit site. Rostral is up.

**Fig. S1.**
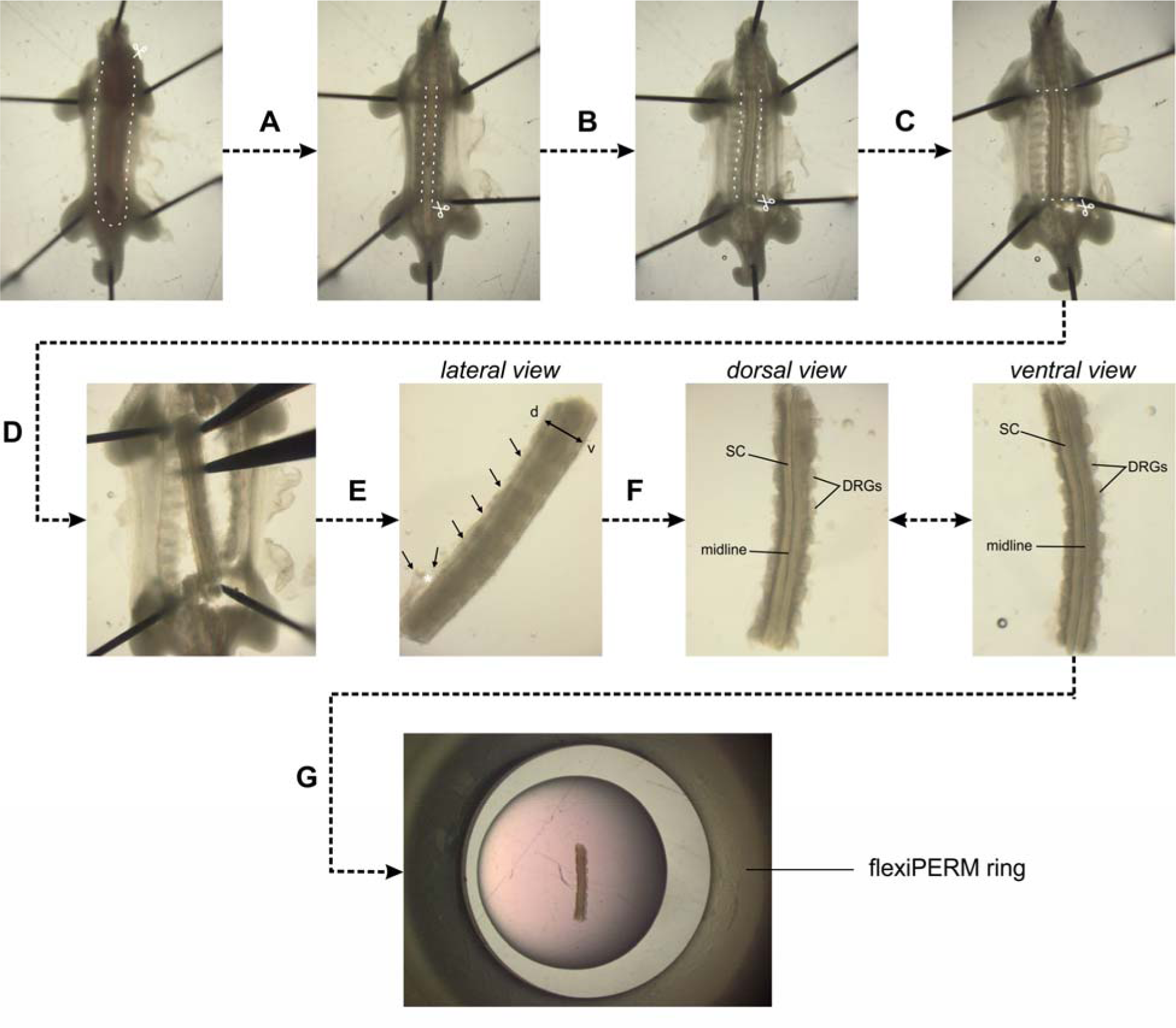
Dissection of intact spinal cords from HH22 chicken embryos. (A) HH22 embryos were pinned down with the dorsal side down in a silicon-coated Petri dish in sterile, cold PBS. Internal organs were removed by first cutting the ventral skin along the dashed lines and pinching out the organs with forceps. (B) Then, a laminectomy was performed, i.e. the ventral vertebrae were cut along the caudal-rostral axis at the level of the outer spinal cord boundaries and the stripe of bone structure was removed with forceps. (C) The visible ventral roots exiting the ventral part of the spinal cord and the peripheral processes of the dorsal root ganglia (DRG) were cut in parallel to the spinal cord without cutting off any DRG. (D) The spinal cord was then cut at the level of the wings and legs. (E) The spinal cord with attached DRG was carefully separated from the rest of the embryo with forceps. Here, special care should be given not to bend the spinal cord by stabilizing the tissue with a second forceps. (F) At this point, the dorsal skin and dermomyotome (black arrows) were removed by first inducing an opening with forceps (white asterisk) taking care not to damage the dorsal spinal cord. Then, using forceps, the dorsal skin and dermomyotome were carefully removed all along the caudal-rostral axis. After this step the dorsal spinal cord should look as clean as the ventral spinal cord with clearly visible midline and no remaining tissues attached (compare dorsal and ventral view). (G) Finally, the intact spinal cord with attached DRG could be embedded as straight as possible in a drop of low-melting agarose-medium mix with the ventral side down. White dashed lines indicate where cuts with small spring scissors should be made. SC, spinal cord; DRG, dorsal root ganglion.

**Fig. S2.**
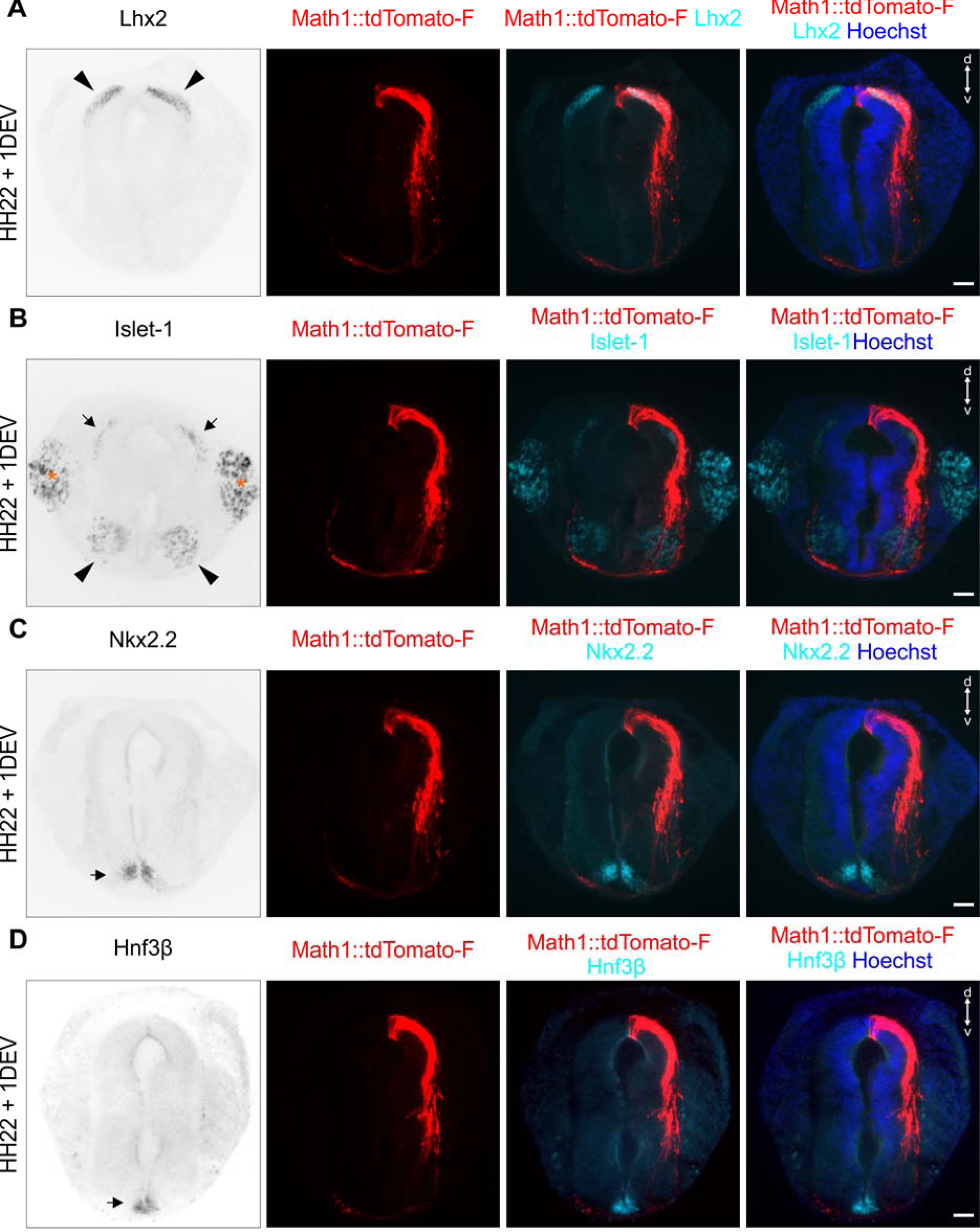
Patterning of cultured intact spinal cords was conserved after one day *ex vivo*. After intact HH22 spinal cords were cultured and imaged for 1 day *ex vivo*, they were fixed and transverse cryosections were immunostained for different dorsal and ventral patterning markers, RFP (Math1-positive dI1 neurons) and counterstained with Hoechst. (A) The dI1 interneuron marker Lhx2 confirmed that these neurons were still localized in the most dorsal part of the spinal cord, as expected (black arrowheads). (B) Islet-1 was used as a marker for DRG neurons (orange asterisks), dI3 interneurons (black arrows) and motoneurons (black arrowheads). All of them maintained the appropriate position: clustered DRG neurons adjacent to the spinal cord; dI3 interneurons localized ventrally of dI1 interneurons; motoneurons on both sides of the ventral spinal cord. (C) Nkx2.2 staining was used to reveal the ventral population of V3 progenitors that are just next to the FP and form the typical inverted V-shape (black arrow). (D) Finally, FP cells forming the intermediate target for dI1 axons were visualized with Hnf3β staining. They were localized at the ventral midline of the spinal cord as expected (black arrow). DEV, day *ex vivo*; d, dorsal; v, ventral. Scale bars: 50 µm.

**Fig. S3.**
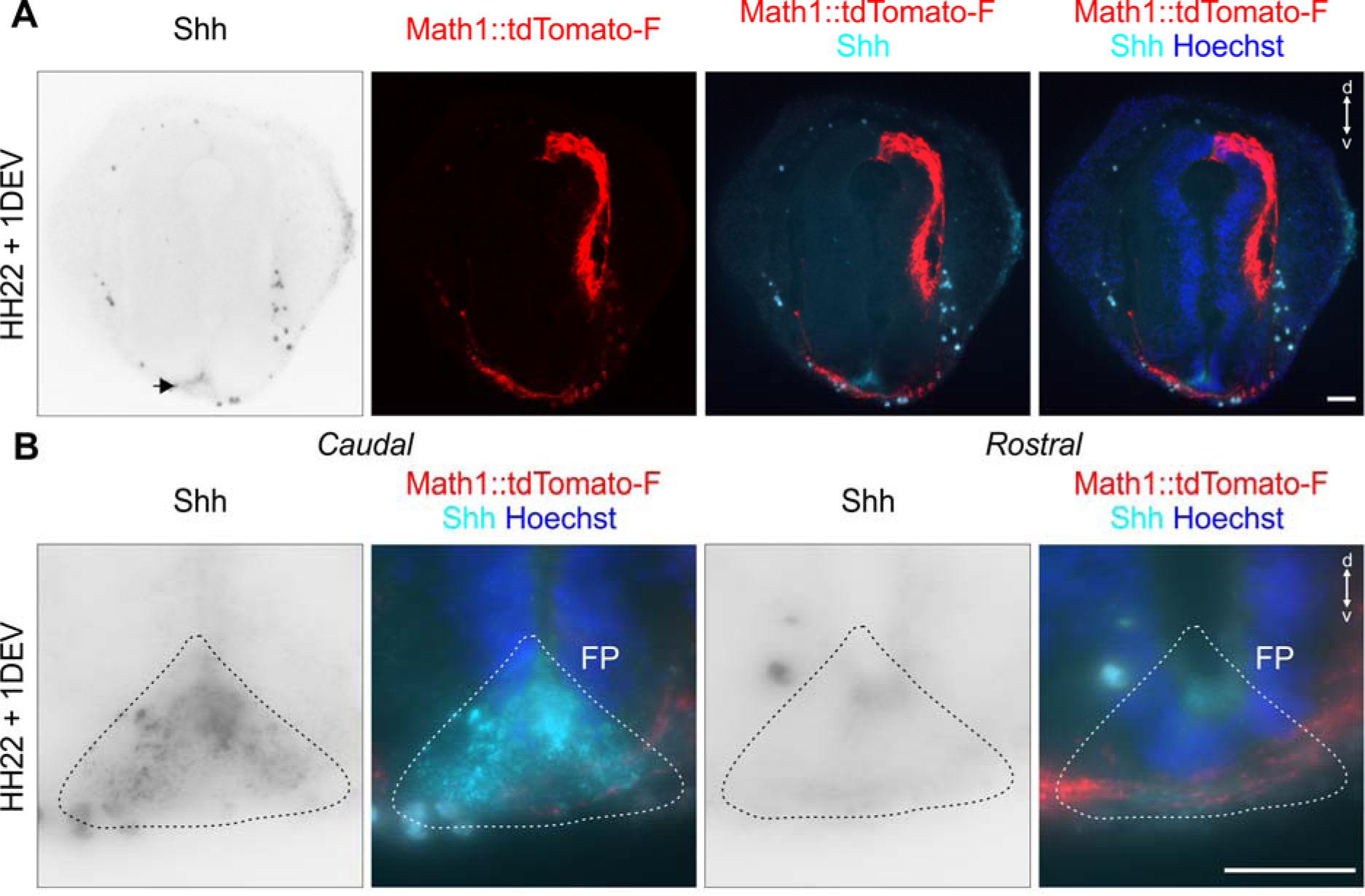
A Shh gradient is still present in a cultured intact spinal cord after one day *ex vivo*. Intact HH22 spinal cords were cultured and imaged for 1 day *ex vivo* before fixation. Transverse cryosections were immunostained with antibodies against Shh (5E1 clone), RFP (Math1-positive dI1 neurons) and counterstained with Hoechst. (A) Shh was still expressed in the FP after one day ex vivo (black arrow). (B) Moreover, in agreement with previous descriptions *in vivo*, Shh was expressed in a decreasing caudal-to-rostral gradient. d, dorsal; v, ventral. Scale bars: 50 µm.

**Fig. S4.**
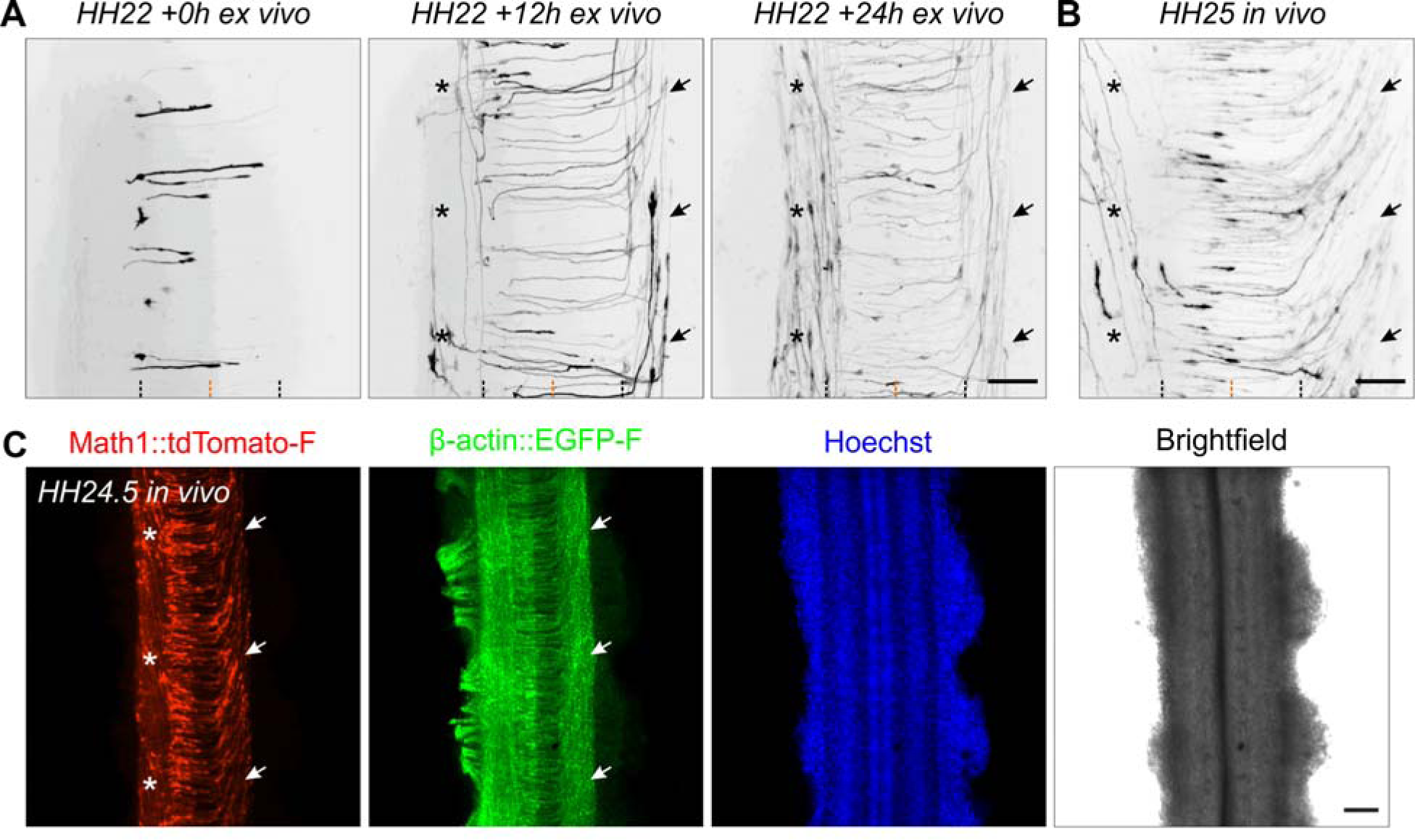
Development of Math1-positive axonal tracts *ex vivo* and *in vivo*. (A) Sequence of 3 images showing dI1 axons crossing the FP after 0, 12 and 24h of culture. After turning rostrally, post-crossing axons started to form the contralateral ventral funiculus (black arrows). After around 12h in culture, a Math1-positive ipsilateral population could be clearly seen in the ipsilateral ventral funiculus (black asterisks). (B) Intact spinal cords dissected at HH25, fixed and mounted similarly to the *ex vivo* culture were imaged the same way. This revealed identical dI1 axonal tracts compared to those seen after 24h of culture of intact spinal cords dissected at HH22, with post-crossing axons forming the contralateral ventral funiculus (black arrows) and ipsilateral axons turning in the ipsilateral ventral funiculus (black asterisks). Black and orange dashed lines represent FP boundaries and midline, respectively. (C) Low magnification overview of an intact HH24.5 spinal cord, fixed, stained for RFP (Math1-positive neurons) and GFP (β-actin transfected cells) and counterstained with Hoechst showing the contralateral ventral funiculus containing post-crossing dI1 axons (white arrows) and the ipsilateral ventral funiculus containing a population of ipsilateral axons (white asterisks). Scale bars: 50 µm (A and B) and 100 µm (C).

**Fig. S5.**
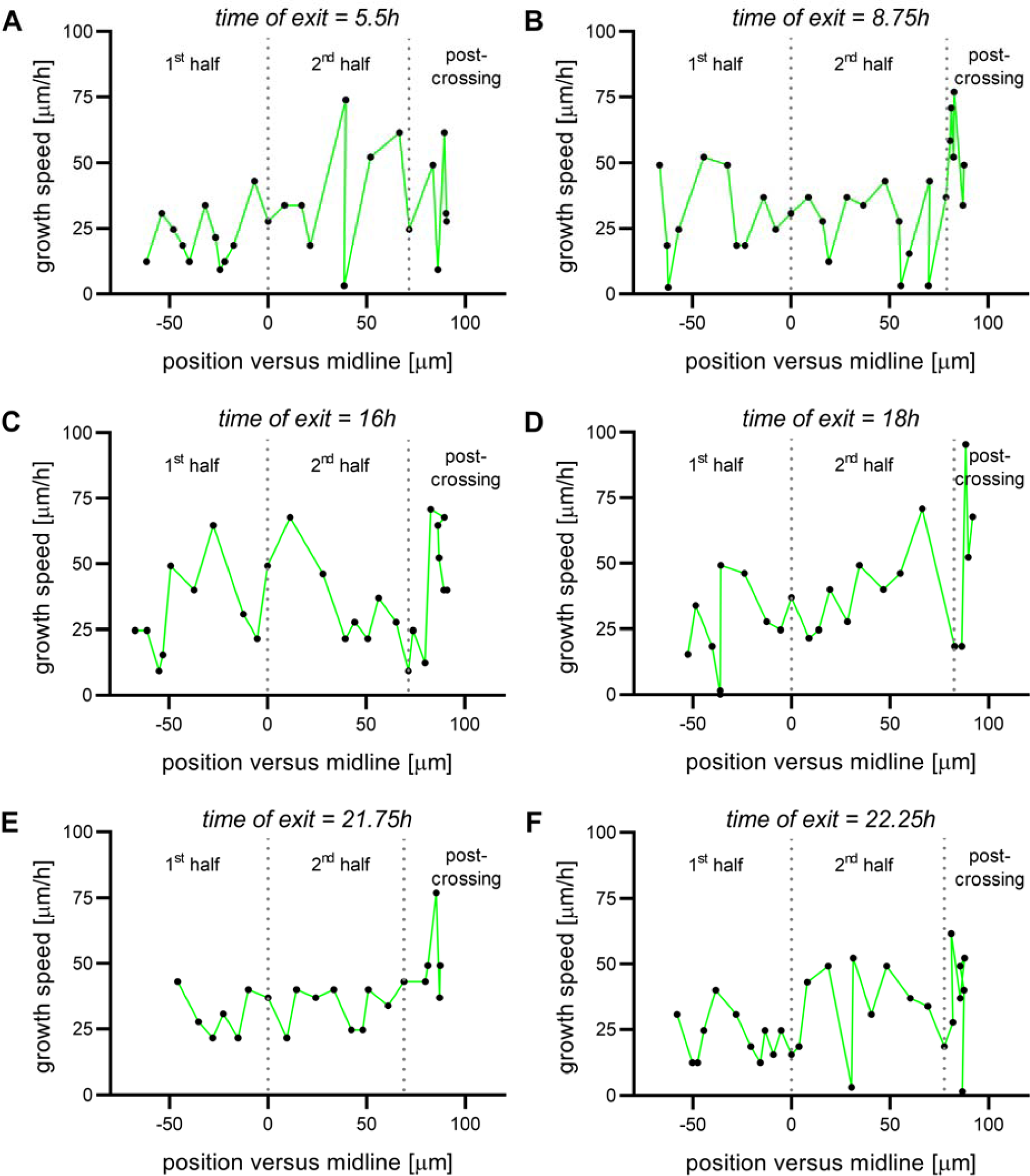
Virtual tracing of axons exiting the FP at different time points. Examples of instantaneous growth speed of axons exiting the FP after 5.5h (A), 8.75h (B), 16h (C), 18h (D), 21.75h (E) or 22.25h (F) of culture that could be extracted and plotted against the position of the growth cone in the FP. Dotted lines represent the time at which the axon crossed the midline or exited the FP. All axons were growing with pulses of acceleration and deceleration. There was no difference between early or late crossing axons.

**Table S1.** Detailed values shown in Figs. 3 and 4.

| **Fig.** | **Name** | **Mean** | **Standard deviation** |  |
| --- | --- | --- | --- | --- |
| Fig.3B | Ent-Mid | 2.81 | 0.96 | n(axons)=298 , N(embryos)=7 |
|  | Mid-Ex | 2.74 | 0.95 | n(axons)=298 , N(embryos)=7 |
|  | Ent-Ex | 5.55 | 1.39 | n(axons)=298 , N(embryos)=7 |
|  | Ex-Turn | 1.39 | 1.03 | n(axons)=298 , N(embryos)=7 |
|  | Ent-Turn | 6.94 | 1.79 | n(axons)=298 , N(embryos)=7 |
| Fig.4B | 1st half | 46.1 | 12.2 | n(growth cones)=127, N(embryos)=7 |
|  | 2nd half | 44.2 | 11 | n(growth cones)=127, N(embryos)=7 |
|  | exit | 105.5 | 25 | n(growth cones)=127, N(embryos)=7 |
|  | after turn | 67.8 | 17.6 | n(growth cones)=127, N(embryos)=7 |
| Fig.4C | 1st half | 41 | 12.6 | n(growth cones)=285, N(embryos)=8 |
|  | 2nd half | 41.8 | 11.5 | n(growth cones)=153, N(embryos)=8 |
|  | exit | 116.9 | 25.7 | n(growth cones)=68, N(embryos)=8 |
|  | after turn | 76.5 | 18.9 | n(growth cones)=102, N(embryos)=8 |

